# Discovery of the Phosphonate Flavophos Produced by *Burkholderia*

**DOI:** 10.64898/2026.01.15.699783

**Authors:** Max A. Simon, Josseline S. Ramos-Figueroa, Vanessa Reyes Lopez, Chayanid Ongpipattanakul, Lingyang Zhu, Constantin Giurgiu, Zoe A. Hoffpauir, Audrey L. Lamb, Satish K. Nair, Wilfred A. van der Donk

## Abstract

Phosphonate natural products have proven value to society as antibiotics and herbicides. They inhibit a range of enzyme targets usually by mimicking the enzyme substrates. In this study, we investigate a family of phosphonate biosynthetic gene clusters (BGCs) found in *Burkholderia*. Heterologous expression in *Escherichia coli* resulted in production of an antimicrobial compound. Spectroscopic characterization and chemical synthesis assigned its structure as 2,4-dioxopentylphosphonic acid. One of the biosynthetic enzymes is a member of the domain of unknown function (DUF) 849 family with homology to β-keto acid cleavage enzymes (BKACEs). In vitro characterization shows that this enzyme catalyzes chemistry that is divergent from previously characterized BKACEs. The observed catalytic activity is explained by a series of co-crystal structures with substrates and intermediates. The BGC also contains a gene encoding lumazine synthase (LS), an essential enzyme in flavin biosynthesis. Biochemical experiments revealed that 2,4-dioxopentylphosphonic acid inhibits LS. In addition, expression of the LS encoded in the BGC, or LS orthologs from a range of organisms, in *E. coli* conferred resistance to the new phosphonate, which we therefore name flavophos.

---

Phosphonic acid natural products have widespread usage in applications from agriculture to medicine.^1–3^ The presence of a hydrolytically stable carbon-phosphorus bond renders phosphonates potent enzyme inhibitors by molecular mimicry of phosphate esters, carboxylic acids, and tetrahedral intermediates in metabolic pathways in all kingdoms of life.^1^ Most known phosphonate biosynthetic pathways derive from the central-metabolite phosphoenolpyruvate (PEP) via formation of the C-P bond by phosphoenolpyruvate phosphomutase (PepM) to generate phosphonopyruvate (PnPy; Figure 1).^4–11^ Thus, bacterial genomes have been mined for the presence of *pepM* genes to discover new phosphonate natural products.^12–14^ The P-C bond forming reaction catalyzed by PepM is thermodynamically unfavorable, with equilibrium favoring PEP by a factor of more than 500.^5^ Phosphonate biosynthesis is usually driven forward by a decarboxylase to form phosphonoacetaldehyde,^9,15–17^ or by an aminotransferase^18,19^ or 2-hydroxyacid dehydrogenase^14,20^ to generate phosphonoalanine and phosphonolactate, respectively.^21^ Less commonly, a homocitrate synthase-like enzyme has been shown to catalyze the condensation of an acetyl group from acetyl-coenzyme A onto PnPy to form 2-phosphonomethylmalate (2-Pmm) (Figure 1).^22,23^ This transformation is encountered in the biosynthesis of FR-900098, fosfomycin, and pantaphos.^22–24^ The fate of 2-Pmm in these pathways differs. During FR-900098 biosynthesis, 2-Pmm undergoes transformations analogous to the citric acid cycle starting with an isomerization (Figure 1).^22^ In the fosfomycin pathway in *Pseudomonas*, 2-Pmm is oxidatively decarboxylated to form 3-oxo-4-phosphonobutanoate (3-OBPn),^25^ and in the pantaphos pathway, 2-Pmm undergoes a dehydration reaction to form phosphonomethylmaleate (PMME) (Figure 1).^26^

**Figure 1.**
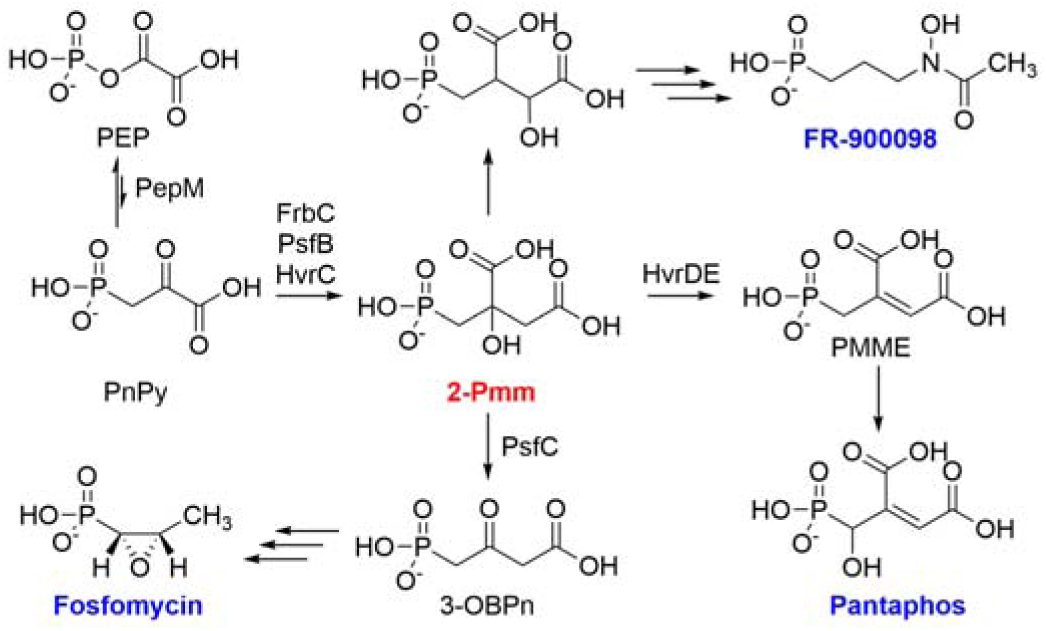
Common steps in the biosynthesis of selected phosphonate natural products. 2-Pmm is generated from phosphonopyruvate by 2-Pmm synthase (FrbC, HvrC or PsfB). This intermediate then undergoes divergent transformations in the biosynthetic pathways to fosfomycin,^23,25^ FR-900098,^22^ and pantaphos.^24^

Prior studies on the biosynthetic pathways towards phosphonic acids focused heavily on organisms within the Actinomycetota phylum,^14^ but many other phyla also possess the machinery to produce phosphonates.^12^ One genus with a surprisingly large number of phosphonate biosynthetic gene clusters (BGCs) per genome is *Burkholderia*, betaproteobacteria within the Pseudomonodota phylum. Analysis of the genomes of diverse strains of this genus^27,28^ revealed BGCs that were predicted to produce new phosphonates. In this study we investigate one of these BGCs (*bsf*) that is widespread in *Burkholderia* and show that the biosynthetic enzymes make a novel phosphonate that we call flavophos. Investigation of its biosynthetic pathway uncovered a new fate for the 3-OBPn intermediate and a new enzymatic reaction for the β-ketoacid cleaving enzyme family. The *bsf* BGC also contains a gene for lumazine synthase (LS) that confers resistance to flavophos, suggesting the new phosphonate likely targets flavin biosynthesis by mimicking the LS substrate 3,4-dihydroxy-2-butanone 4-phosphate (DHBP).

## Results

### Bioinformatic discovery of new phosphonate BGCs involving 2-Pmm

Given the less frequent occurrence of 2-Pmm in phosphonate biosynthetic pathways, we started our genome mining by searching for homologs of the 2-Pmm synthase protein PsfB that is involved in the biosynthesis of fosfomycin in *Pseudomonas*.^23^ The resulting BLAST sequence list was analyzed using the Enzyme Function Initiative Enzyme Similarity Tools (EFI-EST) to generate a sequence similarity network (SSN).^29^ The alignment score threshold was modulated until the sequences of FrbC and PsfB involved in different phosphonate biosynthetic pathways (Figure 1) separated into different groups (Figure 2A). PsfB shares sequence similarity with trans-homocitrate synthase, and therefore a BLAST search using PsfB as the query sequence also returned sequences that were unlikely to be involved in phosphonate biosynthesis if they lacked a nearby *pepM* gene. The full SSN was therefore analyzed by the genomic neighborhood network (GNN) tool of the EFI-EST to generate a list of sequences that co-localized with a gene for a PepM protein (Pfam 13714). Sequence neighborhoods were checked manually to ensure that the BGC did not also encode a phosphoenolpyruvate decarboxylase. The alignment score threshold for the final SSN was adjusted such that sequences whose genome neighborhoods contained nearly identical sets of biosynthetic genes formed distinct groups, suggesting that each corresponded to a biosynthetic gene cluster family (GCF)^30,31^ (Figure 2B).

**Figure 2.**
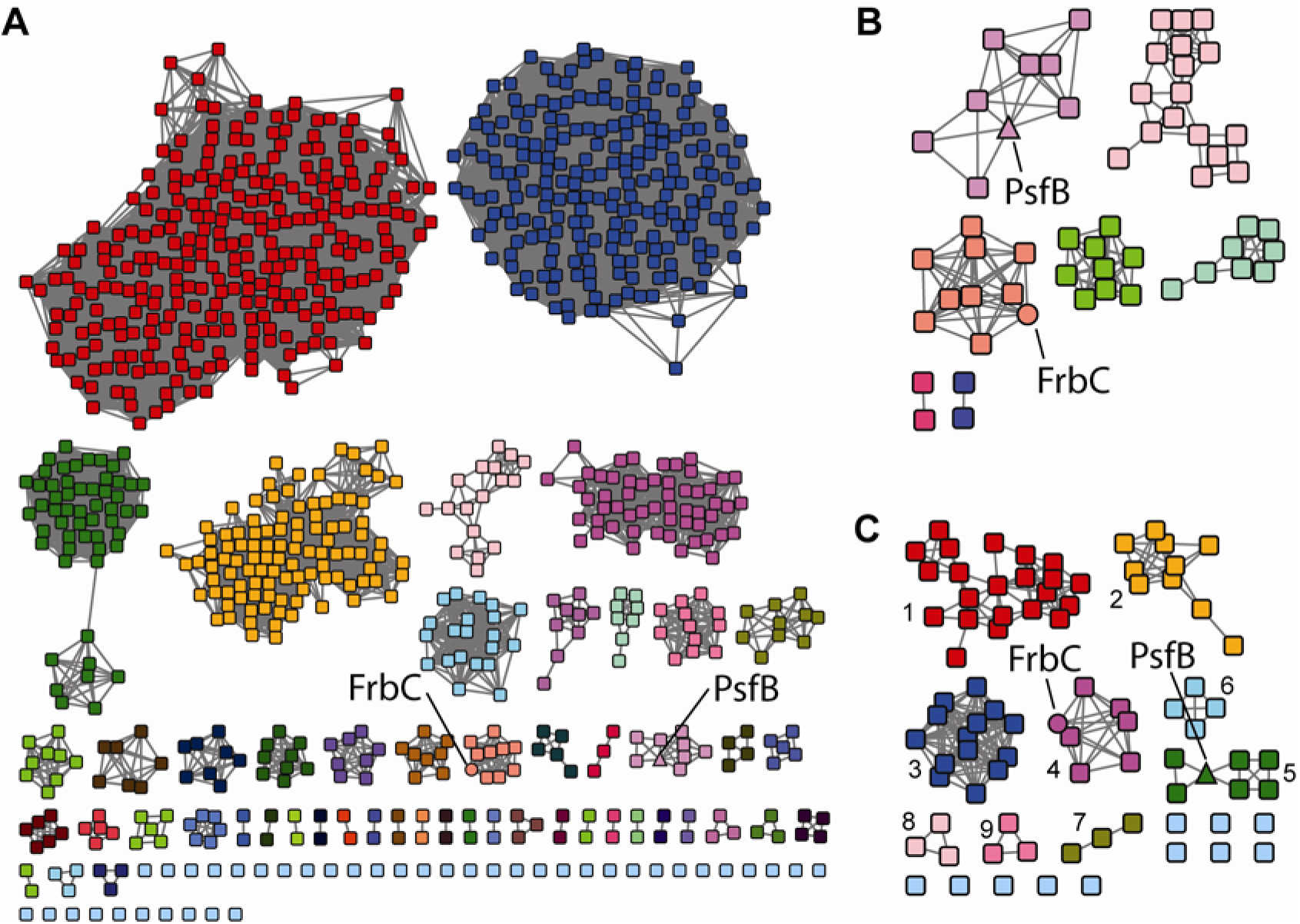
Sequence similarity network of proteins with sequence homology to PsfB. **A**) The sequence of PsfB was used as a BLAST query to identify the top 1000 sequences using the EFI-EST SSN tool.^32^ See Dataset 1 for Cytoscape file. **B**) Phosphonate-related sequences were identified through GNN extraction of sequences that had a PEP mutase Pfam member within 10 ORFs of the PsfB homolog (see Methods and Dataset 2). **C**) The sequences identified in panel B were resubmitted to BLAST analysis, and the retrieved sequences were combined and dereplicated to expand the diversity (Dataset 3). Colors in each panel are independent.

To expand the diversity of sequences, each node from the initial SSN (Figure 2B) was resubmitted to BLAST to return the nearest 1000 sequences. The results for each sequence were combined, de-replicated, and used for SSN and GNN analysis. This procedure increased the diversity of phyla to include Bacillota (Figure 2C; see Cytoscape files provided as Supporting Information datafiles).

The GCFs thus obtained represent several known biosynthetic pathways. GCF 1 (Figure 2C) corresponds to pantaphos biosynthesis,^26^ GCF 4 to FR-900098 biosynthesis,^33^ and GCF 5 to fosfomycin biosynthesis in pseudomonads.^23^ GCFs 2, 3, and 6-9 are predicted to produce unknown products derived from the 2-Pmm intermediate. GCF 3 is restricted to *Burkholderia* species and contains an ortholog of the dinuclear non-heme iron dependent enzyme PsfC described previously that converts 2-Pmm to 3-OBPn during fosfomycin biosynthesis by GCF 5 (Figure 1).^25^ Thus, GCFs 3 and 5 share the enzymatic steps leading to the intermediate 3-OBPn (Figure 3). However, the remaining genes in GCF 3 differ from those in the fosfomycin pathway (see next section) suggesting that the BGCs in GCF 3 direct the biosynthesis of a distinct phosphonate natural product.

**Figure 3.**
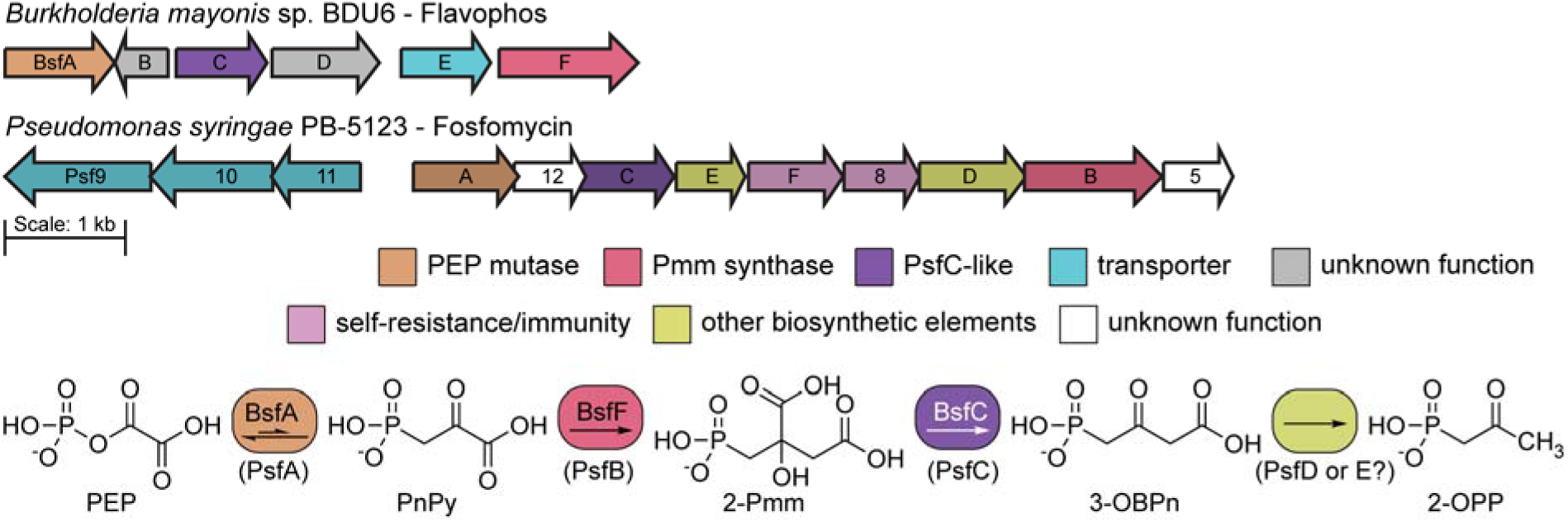
A phosphonic acid biosynthetic pathway found in *Burkholderia* proceeds through the 3-OBPn intermediate. Representative biosynthetic gene clusters for GCF 3 (top) and GCF 5 (bottom) that are predicted to generate the 2-Pmm intermediate. Enzymes predicted to catalyze identical chemical steps are in the same color. The *B. mayonis* sp. BDU6 BGC shares three genes with the fosfomycin BGC from *P. syringae* PB-5123. An anticipated partial pathway is drawn using these three gene products, ending with the BsfC/PsfC product, 3-OBPn. In the fosfomycin pathway 3-OBPn is decarboxylated to 2-oxopropylphosphonate (2-OPP), possibly by one of the currently uncharacterized enzymes encoded in the BGC.

### The *bsf* phosphonate biosynthetic gene cluster in *Burkholderia*

The phosphonate biosynthetic genes found within GCF 3 are highly conserved and consist of six genes. The BGC in the type strain *Burkholderia mayonis* sp. BDU6 that is the focus of this study,^34^ encodes homologs of PepM (BsfA; accession AOJ04873.1), Pmm synthase (BsfF; AOJ04877.1), and PsfC^25^ (BsfC; AOJ04874.1) (Figure 3, Table S1).^25^ The synteny of these genes in GCF 3 is different from that of members of GCF 5, one of which was characterized for the *Pseudomonas* fosfomycin pathway (Figure 3). Nevertheless, a predicted partial biosynthetic pathway can be drawn based on the anticipated activity of the enzymes (Figure 3). The remaining genes in GCF 3 are predicted to encode a β-ketoacid cleavage enzyme (BsfD; AOJ04875.1),^35,36^ and a multidrug transporter of the EamA family (BsfE; AOJ04876.1) (Figure 3).^37^ In other pathways, β-ketoacid cleavage enzymes (BKACEs) take β-ketoacids and acetyl-coenzyme A (CoA) and convert them into acetoacetate and a CoA-thioester product of the β-keto acid that is shortened by two carbons (Figure S1).^35,36,38^ The only gene found on the reverse strand for GCF 3 appears to encode a homolog to lumazine synthase (BsfB; AOJ05795.1), an essential enzyme in riboflavin biosynthesis (Figure 3).^39^ While *bsfB* may also play a role in the biosynthesis of the *bsf* product, its presence in the opposite reading frame suggests the gene may instead have a role as an immunity gene, protecting the producing organism from any toxic effects of the final natural product.^40,41^ Of the remaining unknown genes within the presumed biosynthetic cluster boundaries, *bsfE* may act as a transporter for the final natural product.

### BsfD converts 3-OBPn to a new phosphonate with bioactivity

To establish the role of BsfD, we co-expressed the protein in *E. coli* together with PsfABC from *Pseudomonas syringae* PB-5123, which generates 3-OBPn along with the decarboxylated break-down product 2-oxopropylphosphonate (2-OPP).^25^ Codon-optimized genes encoding BsfD were purchased that were identified in two strains harboring the *bsf* BGC in their genomes, *B. mayonis* sp. BDU6 (*Bm*BsfD) and *Burkholderia stagnalis* (*Bs*BsfD). The genes were cloned into the multiple-cloning site (MCS) 2 of a pETDuet vector containing the *psfC* gene in MCS1.^25^ Co-expression of *psfABC* with *bsfD* from either *Burkholderia* strain in *E. coli* resulted in new peaks in the ^31^P NMR spectrum of the lysate, along with the expected peaks for 2-Pmm, 3-OBPn and 2-OPP (Figure 4C). The lysate was also assayed against a bacterial indicator strain, *E. coli* WM6242 λ(DE3), showing a clear zone of growth inhibition (Figure 4D). This engineered indicator strain has the endogenous phosphonate transporter operon (*phn*) under the control of the inducible Tac promoter.^22^ Upon induction of expression of the operon, the indicator strain is highly sensitive to phosphonates overcoming the usually low cell permeability of these compounds. All four biosynthetic genes were required for bioactivity of the phosphonate product of the *bsf* BGC, as removal of any gene resulted in loss of observed bioactivity.

**Figure 4.**
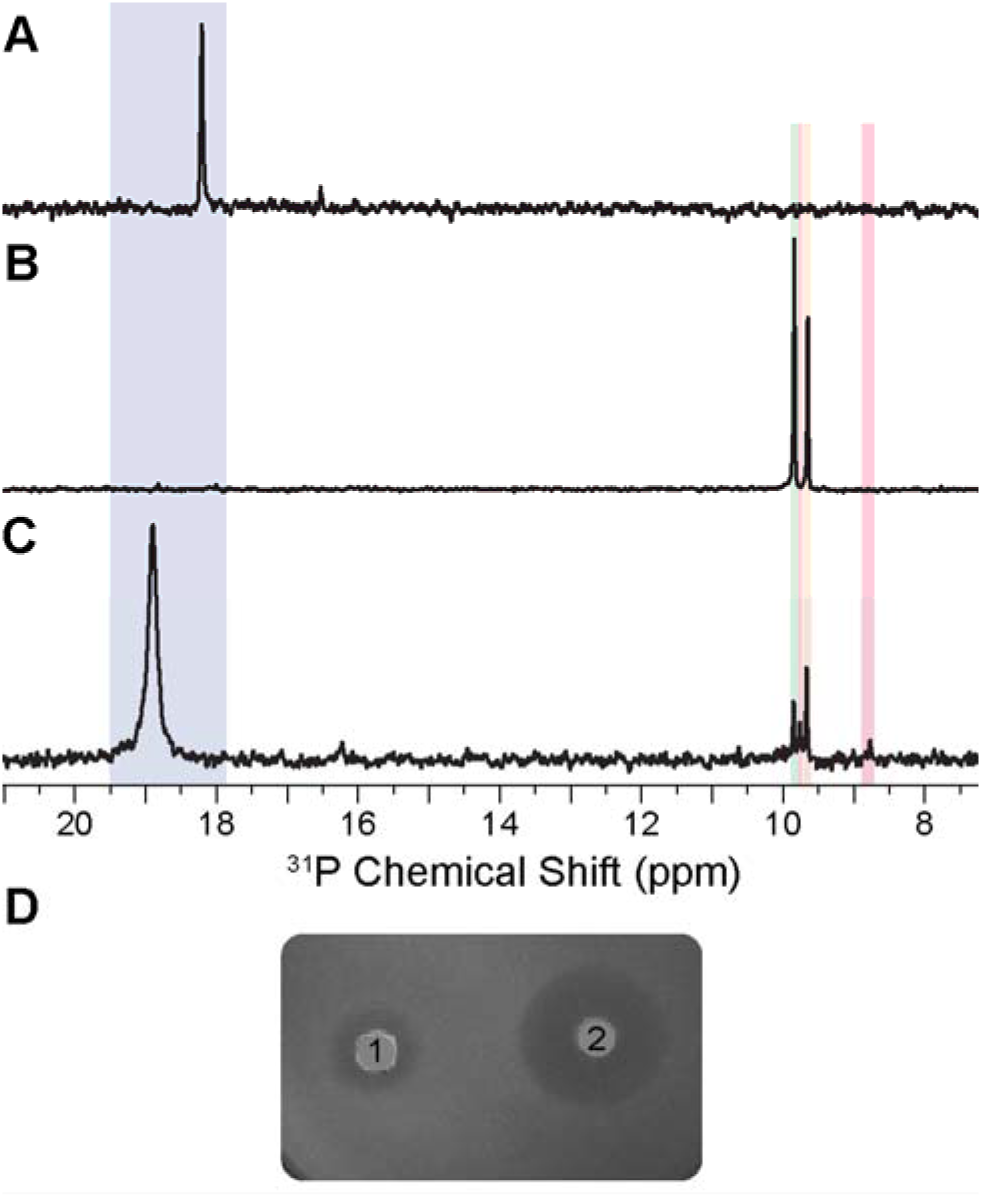
Heterologous expression of PsfABC + BsfD in *E. coli* and bioactivity screening of the cell lysates. ^31^P NMR analysis of *E. coli* cell lysates that co-expressed **A**) *psfAB*, **B**) *psfABC*, and **C**) *psfABC* with *Bm-bsfD*. 2-Pmm (blue, ∼18 ppm), 3-OBPn (orange, 9.7 ppm), and 2-OPP (green, 9.9 ppm) are observed. A new product is generated in sample **C** highlighted in red at 8.8 ppm. ^31^P chemical shifts of phosphonates are highly sensitive to pH near their pK_a_ resulting in small shifts from experiment to experiment. The identity of the peaks was therefore confirmed by spiking with authentic standards. **D**) Bioactivity screening of cell lysates expressing *psfABC* and *bsfD* against *E. coli* WM6242 λ(DE3) plated on LB-agar containing IPTG to induce expression of the *phn* operon. 1) Lysate from cells expressing *psfABC* and *Bm-bsfD*, 2) cell lysate from cells expressing *psfABC* and *Bs-bsfD*.

### Reconstitution of flavophos generation *in vitro*

Despite many different attempts, we were unable to purify the bioactive product from heterologous expression in *E. coli*. We therefore explored *in vitro* enzymatic assays to access the bioactive product. *Bs*BsfD and *Bm*BsfD were expressed and isolated as N-terminal His_6_-fusion proteins as described previously for other members of the BKACE family of proteins.^36^ The inclusion of sorbitol (0.5 M) and betaine (5 mM) was necessary for soluble protein isolation.^38^ The expression of *Bs*BsfD resulted in higher yield and therefore all in vitro experiments were performed with this ortholog. BKACEs utilize AcCoA as a co-substrate with β-ketoacid-containing compounds to generate CoA-esters and acetoacetate (Figure S1).^35,38,42^ We therefore predicted that BsfD would generate phosphonoacetyl-CoA, with acetoacetate as a by-product (Figure S1).

To probe whether BsfD performs the anticipated reaction, we used ^13^C-labeled AcCoA (2-^13^C-AcCoA) as a co-substrate, and monitored production of any 4-^13^C-acetoacetate by ^13^C NMR spectroscopy. A new peak corresponding to a ^13^C-labeled methyl group at 27.3 ppm was produced only when all reaction components were included (Figure S2). However, when this sample was supplemented with a standard of acetoacetate, a new resonance appeared upfield at 26.3 ppm (Figure S2) indicating that the enzymatic product is not that expected based on BKACE enzymology. Additional spiking of this sample with acetone, a potential degradation product of acetoacetate, showed that this compound is also not giving rise to the new peak observed in the co-expression sample (Figure S2).

Further NMR experiments utilizing the material derived from the in vitro reaction with ^13^C-AcCoA allowed partial structural characterization of the phosphonate product of BsfD using a combination of ^1^H-^13^C HMBC and HSQC spectroscopy (Figure 5). The data show a terminal methyl group as well as a pair of methylene protons, both adjacent to the same carbonyl. One of the methylene groups displays a doublet from coupling to ^31^P (^2^*J*_PH_ = 21 Hz). The collective data suggested that the new phosphonate is 2,4-dioxopentyl phosphonic acid (Figure 5E). This compound was therefore prepared from diethyl-protected 2-oxopropylphosphonate and ethyl acetate,^43^ followed by deprotection and purification.^44^ Synthetic 2,4-dioxopentyl phosphonic acid displayed identical NMR and MS features as the product from the *bsf* BGC.^44^ Addition of the authentic standard to an *in vitro* BsfD reaction resulted in an increase in the ^31^P resonance of enzymatic product, confirming that the product generated in vitro and in *E. coli* (flavophos) is 2,4-dioxopentylphosphonic acid (Figure 5CD, and Figure S3). Bioactivity testing of the synthetic material also demonstrated the same antimicrobial phenotype as observed with *in vivo* and *in vitro* prepared flavophos.

**Figure 5.**
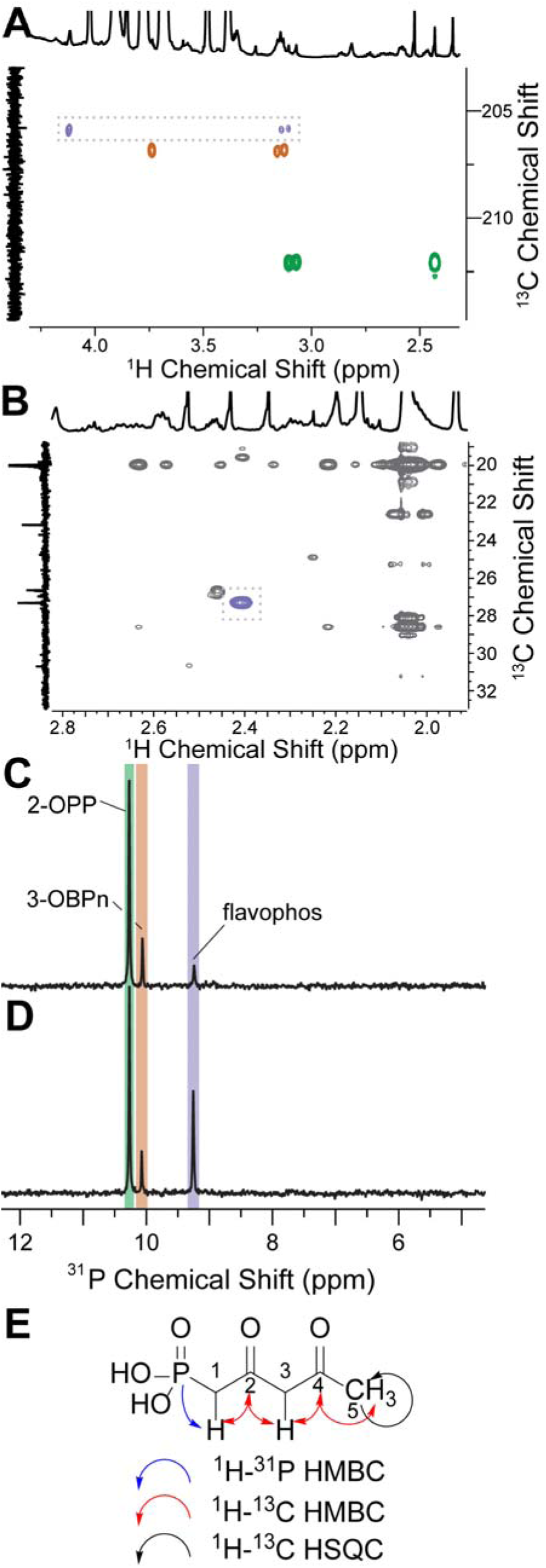
Preliminary structural elucidation of the *bsf* product and spiking with synthetic material. **A**) ^1^H-^13^C HMBC NMR spectrum of the *in vitro* BsfD reaction producing flavophos. Cross-peaks associated with 2-OPP are highlighted in green, with 3-OBPn in orange, and with flavophos in purple (dashed box). A doublet is observed at ∼ 3.2 ppm in the ^1^H dimension with a ^2^*J*_PH_ value of 21 Hz. These protons couple to a carbonyl carbon at 206 ppm (purple). The same carbonyl displays a crosspeak to an additional set of protons at 4.2 ppm. Integration of these two sets of protons results in a 1:1 ratio. **B**) ^1^H-^13^C HSQC NMR spectrum illustrating coupling of the ^13^C-labeled methyl group in the product (27 ppm) with a ^1^H signal at 2.4 ppm (highlighted in purple). **C**) *In vitro* incubation of purified *Bs*BsfD with 2-^13^C-AcCoA and the lysate of cells co-expressing *psfABC* that produce 3-OBPn and 2-OPP. The assay results in a single new product by ^31^P NMR spectroscopy (highlighted in purple) at 9.2 ppm. **D**) Addition of synthetic 2,4-dioxopentylphosphonic acid shows an increase in the signal associated with flavophos. **E**) Structure of flavophos and the correlations in NMR experiments underlying its structural assignment.

### Structure and mechanism of BsfD

Sequence analysis shows BsfD to fall within the DUF849 family of enzymes that are functionally diverse,^35^ with the reaction catalyzed by BsfD representing new chemistry. To obtain further insights into the reaction mechanism, we carried out crystallization of *Bs*BsfD and determined its structures in three different states: 1) in the absence of any ligands (apo form, 1.72 Å resolution, Figure 6A), 2) with bound coenzyme A (CoA form, 1.5 Å resolution, Figure 6B), and 3) from crystals grown in the presence of acetyl-CoA and 3-OBPn that yielded structures with bound CoA and either substrate (short soak, 1.81 Å resolution, Figure 6C) or the presumptive product (long soak, 1.74 Å resolution, Figure 6D). Initial crystallographic phases were determined by the molecular replacement method using the coordinates of a putative 3-keto-5-aminohexanoate cleavage enzyme (Kce; Figure S1) from *Ralstonia eutropha* (*Ra*Kce) as a search probe (PDB code: 3C6C, 40% sequence identity).

**Figure 6.**
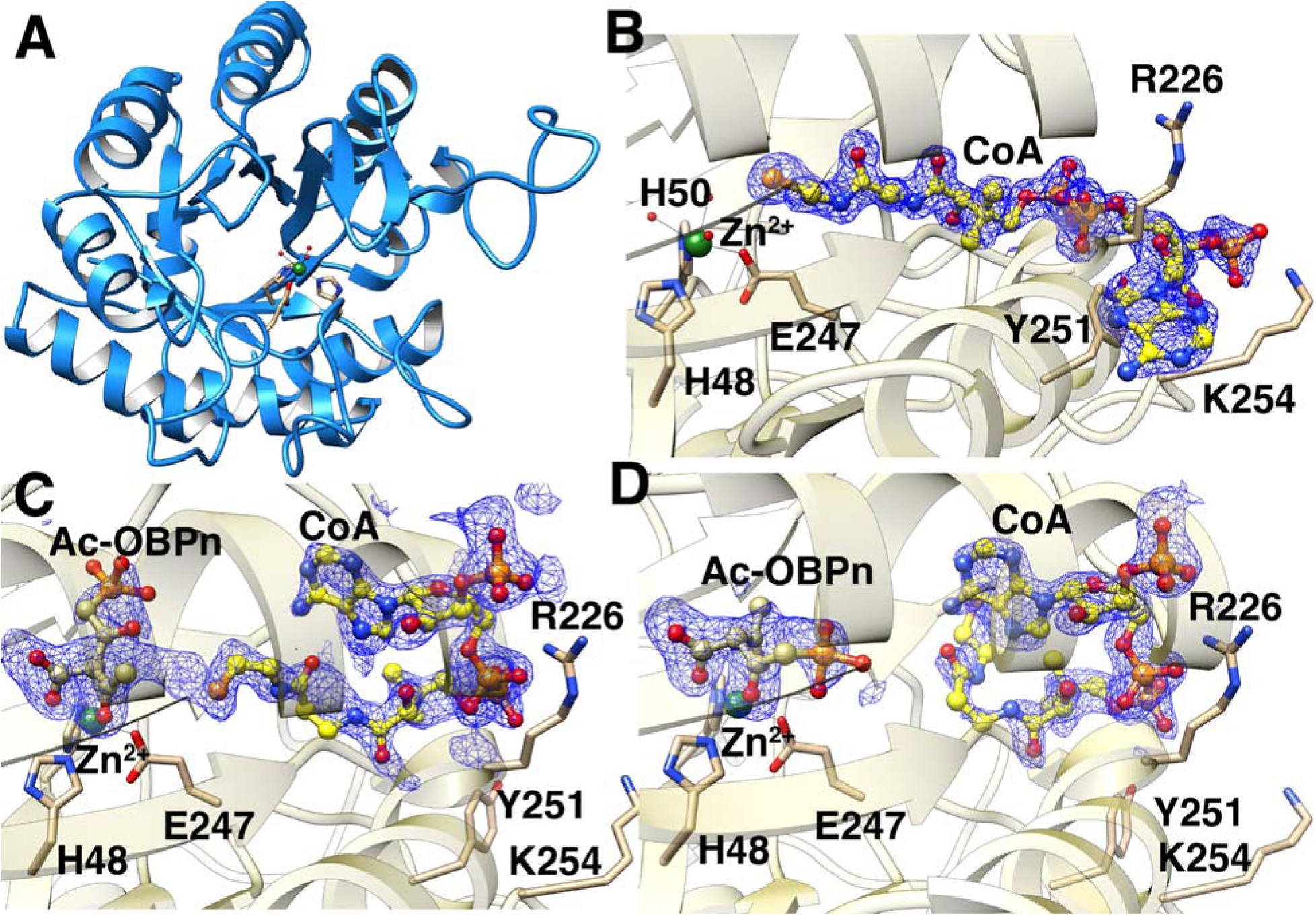
Crystal structures of *Bs*BsfD. **A**) The overall structure of *Bs*BsfD with a Zn^2+^ ion (green sphere) in the active site (PDB ID 9ZSC). **B**) Co-crystal structure of BsfD with bound coenzyme A (PDB ID 9ZSH). Interacting residues and Zn^2+^ ligands are displayed. **C**) Co-crystal structure of BsfD incubated with AcCoA and 3-OBPn for six hours showing electron density consistent with the intermediate Ac-OBPn and HSCoA (PDB ID 9ZSV) in an arrangement expected for the products immediately after catalysis (Figure 7). **D**) Co-crystal structure of BsfD incubated for two days with AcCoA and 3-OBPn showing electron density consistent with rearranged HSCoA and the product Ac-OBPn (PDB ID 9ZSN).

The overall structure of *Bs*BsfD (Figure 6A, Table S2) recapitulates the (β/α)_8_ TIM barrel fold observed in the structures of *Ra*Kce, consistent with the sequence similarity between these enzymes (Z-score=43.6, RMSD of 1.4 Å over 288 aligned Cα residues). A search using the DALI server^45^ against the Protein Data Bank shows similar levels of fold conservation with other members of the DUF849 protein family from bacteria and archaea, whose structures were determined by various structural genomic consortia (PDB codes: 2Y7F, 3LOT, 3NO5, 8RIP) all with Z-score values in the range of 38-40 and with mean Cα RMSD values around 1.5 Å. Lower structural similarity (Z-score values of 15-30) was noted with other enzymes that contain a TIM barrel fold. A Zn^2+^ ion is located at the center of the TIM barrel of BsfD where the ion is engaged by protein residues His48, His50 and Glu247 (Figure 6B, S4). These residues and three solvent molecules complete an octahedral coordination geometry at the metal. The identity of the metal was confirmed by calculation of anomalous difference Fourier map using data collected at the zinc absorption edge (Figure S5). Difference maps calculated with Bijvoet pairs showed strong peaks at the location of the zinc.

The structure of *Bs*BsfD with CoA (Figure 6B) shows that the cofactor occupies a binding mode like that observed in the structures of the *Pd*Kce from *Paracoccus denitrificans* (PDB code: 8RIP; Figure S1B),^46^ and in that of the quorum signal biosynthetic enzyme ObcA, which catalyzes a metal- and AcCoA-dependent step during the breakdown of oxaloacetate to oxalic acid and acetoacetate (PDB code: 4NNC; Figure S1C).^47^ ObcA is not a member of the DUF849 fold as the protein contains an N-terminal extension of ∼200 residues that provides several residues that interact with and fix the position of the bound CoA. In contrast, the co-crystal structure of *Bs*BsfD shows limited interactions with CoA, with the 3’-phosphoadenosine moiety engaged by only Arg226 and Lys254 above and below the plane of the ring (Figure 6B), with the latter residue also interacting with the 3’-phosphate. Such limited interactions with the 3’-phosphoadenosine are also observed in the structure of the *Pd*Kce.^46^ In this structure (as well as the other structures), Tyr251 is positioned to sterically direct the CoA pantetheine group away from solvent and inwards to the active site of the enzyme.

To obtain structures with bound substrate, we enzymatically generated 3-OBPn in *E. coli* using PsfABC and rapidly isolated the small molecule products (including the break-down product 2-OPP) using an Amicon concentrator with a 10 kDa molecular weight cut off to remove the proteins. The mixture was incubated with *Bs*BsfD and acetyl-CoA and subjected to co-crystallization efforts. Diffraction quality crystals were subsequently incubated with additional aliquots of 3-OBPn for short and long time increments before collection of diffraction data. Even though the ligand composition used for crystallization contained both intact 3-OBPn and the decarboxylated break-down product 2-OPP, electron density observed in either of the resultant structures is consistent with enzyme bound intermediates that have not decarboxylated and could be confidently modeled owing to the high resolution of the structures.

The co-crystals with both substrates occupy a different space-group with two copies in the asymmetric unit allowing for multiple independent and unbiased views of the bound ligands. Soaking of the AcCoA crystals with 3-OBPn for six hours yielded a structure with density for CoA directed into the active site directly adjacent to the acetylated phosphonate (Ac-OBPn, Figure 6C and 7A) (1.81 Å resolution), while soaking for two days yielded a structure (1.74 Å resolution) with Ac-OBPn near the Zn^2+^ but with the CoA phosphopantetheine positioned away from the active site (LigPlots are provided as Figures S6 and S7). The short soak structure shows compelling evidence for a covalent adduct between acetyl-CoA and 3-OBPn, but was not modelled as such due to the mixture of likely products formed in situ. The most notable feature of each structure is the orientation of the 3’-phosphoadenosine group of CoA, which is flipped towards the direction of the active site (Figure 6D and S7), effectively occluding the bound substrates within a closed cavity. The movement of the phosphoadenosine also directs the phosphopantetheine away from the metal center, which should allow for product release after acetyl transfer. A comparison of the structure with 3-OBPn soaked for six hours versus that for two days shows that the Ac-OBPn undergoes a rearrangement in the active site after the CoA phosphopantetheine moves away to position the C4 of the product acid (for atom numbering, see Figure 7) adjacent to the Zn^2+^ (Figure S6). Furthermore the carbonyl oxygens at C2 and C4’ coordinate to the Zn^2+^ and will help stabilize the developing negative charge on these oxygens during decarboxylation^48^ to the final product flavophos (Figure 6D and S6). A comparison of the structures of *CCa*Kce from *Candidatus* Cloacimonas acidaminovorans in the absence of CoA (or the acetyl-thioester) bound to the substrate 3-keto-amino-hexanoate (PDB Code: 2Y7F) or with bound product acetoacetate (PDB Code:2Y7G) shows a similar flipping of the acetoacetate product at the metal center.^38^ In contrast, co-crystal structures of *Pd*Kce with CoA and substrate or product (Figure S1) show only a single conformer in which the carboxyl groups of acetoacetate or malonate were stabilized by the metal ion.^46^

**Figure 7.**
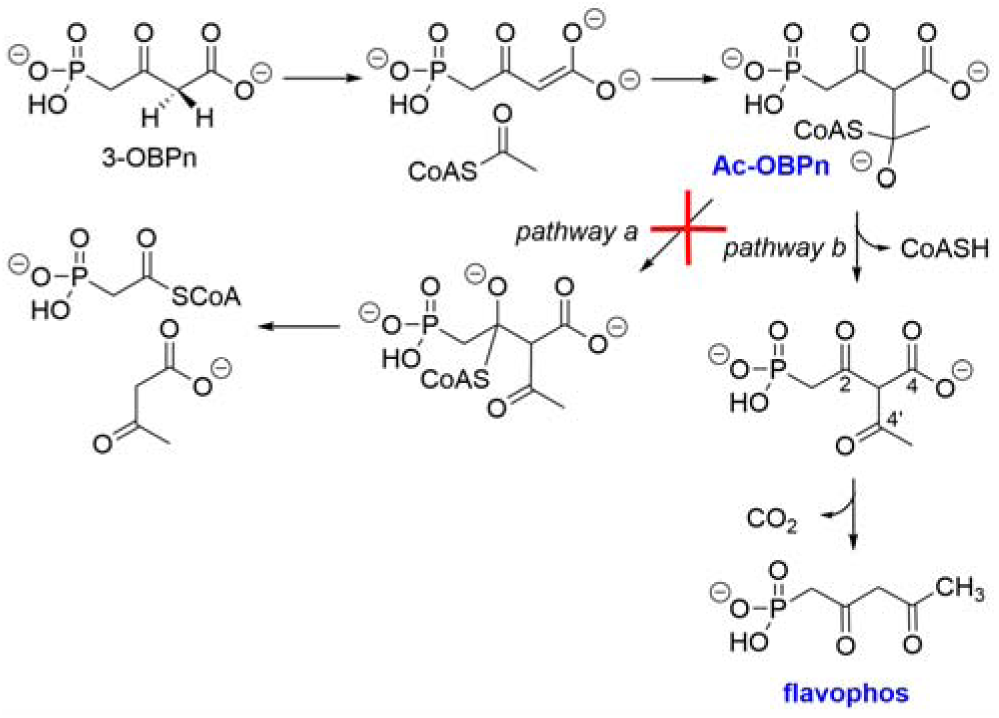
Reaction mechanism of a canonical BKACE reaction (pathway a) and the proposed mechanism for the observed reaction with BsfD (pathway b).

A comparison of the structures of *Bs*BsfD with that of the biochemically and structurally characterized β-ketoacid cleavage enzymes *Cca*Kce (PDB code: 2Y7F),^38^ *Pd*Kce (PDB code: 8RIP),^46^ and the related enzyme ObcA (PDB code: 4NNC)^47^ suggests differences that may distinguish the mechanism of *Bs*BsfD from the reactions catalyzed by BKACEs. First, these three BKACE sequences contain a Tyr residue (Tyr173 in *Pd*Kce and Tyr145 in *CCa*Kce) that is thought to be involved in catalysis by interacting with the sulfur in CoA;^38^ in ObcA (Figure S1C), Tyr322 occupies a spatially equivalent position. Mutation of Tyr322 in ObcA resulted in a loss of activity.^47^ Although this residue has been implicated in canonical BKACE catalysis its exact function is unclear. In *Bs*BsfD (but also in *Ra*Kce), the equivalent residue is Phe166 (Figure S4). Because the reaction catalyzed by *Ra*Kce has not been experimentally determined, we investigated whether replacement of Phe166 with Tyr in BsfD might catalyze the canonical BKACE reaction and generate phosphonoacetyl-CoA (Figure S1D). Unlike WT BsfD, co-expression of BsfD-F166Y with PsfABC in *E. coli* did not result in a peak for flavophos in the ^31^P NMR spectrum of the lysate and only 3-OBPn and 2-OPP were observed. We tried to purify BsfD-F166Y using identical procedures as for WT BsfD, but were unable to obtain soluble protein. Hence, we were unable to establish whether the replacement of Tyr166 with Phe in BsfD is related to the observed change in chemistry compared to canonical BKACEs.

Second, the flipping of the CoA 3’-phosphoadenosine observed in our structures positions the thiolate away from the ligand and disfavors formation of the adduct of CoASH to C2 of acetyl-OBPn that would be necessary for cleavage of the C2-C3 bond in a retro-Claisen reaction catalyzed by BKACEs (Figure 7, pathway a). An equivalent shift of the 3’-phosphoadenosine is disabled by secondary structural elements in canonical BKACEs. For example, the structure of *Pd*Kce contains two α-helices that cap the active site and would restrict any movement of the CoA.

The structure also helps provide a rationale for the preference of *Bs*BsfD for a phosphonate substrate. Notably, the sequence of *Bs*BsfD contains an insertion of ∼20 residues in the loop following strand β3 (Figure S4), which positions a unique basic residue (Arg91) near the phosphonate. In other characterized BKACEs, this loop is either shortened or a basic residue is not present in an equivalent position (Figure S4). In the *Bs*BsfD structures, Arg91 is positioned to stabilize the phosphonate only after the molecule flips following acetyl transfer and may play a role in product release or in the flipping process.

### The target of flavophos is lumazine synthase in riboflavin biosynthesis

BsfB, the duplicate copy of lumazine synthase (LS) encoded within the flavophos BGC, is not necessary for flavophos production. BsfB contains all of the conserved residues that are important for the catalytic function of lumazine synthases (Phe22, His88, Arg127).^49,50^ Many natural product BGCs contain self-immunity genes that are duplicate versions of house-keeping genes that encode a resistant form of the protein to protect the producing organisms.^40,51^ We therefore investigated if BsfB would confer resistance to flavophos when expressed in *E. coli* WM6242 λ(DE3). Indeed, expression of *Bm*BsfB in *E. coli* WM6242 λ(DE3) using an arabinose inducible promoter in the plasmid pBAD^52^ resulted in abolishment of flavophos activity (Figure S8A). To probe whether BsfB was unique in its resistance properties, the house-keeping forms of LS in *E. coli* (*Ec*LS; 33% sequence identity) and *B. subtilis* (*Bsu*LS) were cloned into the same pBAD vector and individually overexpressed in the indicator strain, again resulting in elimination of antibacterial activity (Figure S8B).^41^ Similarly, LS from *Aquifex aeolicus,* a thermophile with an optimum growth temperature of around 90 °C,^53^ conferred resistance (Figure S8B). These findings suggest that BsfB is not unique as a resistance marker.

Lumazine synthases have been studied extensively for their biotechnological applications. LSs from various different organisms including *E. coli, B. subtilis*, and *A. aeolicus* form a highly stable capsid that is typically composed of a dodecamer of pentamers (60-mer).^50,53,54^ Encapsulation of biosynthetic intermediates in this structure is thought to enhance the biosynthetic output of the pathway.^55^ The size, shape, and composition of these capsids, as well as their caged components vary between species,^50^ and has been the focus of protein engineering studies due to their encapsulation properties.^56–61^ Encapsulation of flavophos could therefore be a resistance mechanism that does not rely on the biosynthetic activity of LS. The observation that hyperthermophilic LS from *A. aeolicus* confers resistance against flavophos to *E. coli* (Figure S8B), even though its activity at the growth temperature used (37 °C) would be very low, supports sequestration as a mechanism of resistance. We therefore also evaluated the LS from *Mycobacterium tuberculosis* (*Mt*LS; RibH), which does not form a protein cage but assembles into homopentamers.^62^ Expression of *Mt*LS in *E. coli* WM6242 λ(DE3) indeed also conferred resistance to flavophos (Figure S9A). These observations hinted that the mechanism of action of flavophos involves inhibition of the catalytic activity of LS that can be overcome by overexpression of LS.

To test the hypothesis that the resistance requires enzymatic activity of the LS, we also investigated two mutants of LS from *B. subtilis* that were demonstrated in previous studies to have very low activities, R127T and the H88K/K131N double mutant.^63^ Expression of both mutants from the arabinose inducible pBAD vector resulted in resistance against flavophos activity (Figure S9B). Collectively, these findings suggest that the observed resistance by expression of LS may be achieved both by sequestration of flavophos or by overcoming inhibition of LS activity.

In an attempt to obtain further evidence for LS being the target of flavophos, we also generated spontaneous resistance mutants of *E. coli* WM6242 λ(DE3) and sequenced the genomes of four mutants. No mutations were found in the gene for the LS enzyme or in the surrounding non-coding regions. Instead, every resistant mutant had one or more mutations in the *phn* operon that facilitates phosphonate uptake in *E. coli* WM6242 λ(DE3) (Table S3). We selected four more spontaneous resistance mutants and amplified and sequenced their *phn* operons. All contained mutations in their *phn* operons that would be expected to inactivate the phosphonate uptake process. Hence, resistance against flavophos readily arises by mutations of the phosphonate transporter system under the conditions where this transporter is not essential. Thus, spontaneous resistance mutants could not be used to provide further support for LS as the molecular target. Media supplementation with excess riboflavin only slightly reduced the antibacterial activity of flavophos. This observation was expected because most bacteria, including *E. coli*, do not contain a riboflavin transporter, and therefore cannot take up riboflavin from their environment.^64,65^

## Mechanism of LS inhibition

The structure of flavophos resembles one of the substrates of LS and could form structures that mimic intermediates in the reaction mechanism proposed for lumazine synthase (Figure 8A).^39,49,63^ Reaction of 3,4-dihydroxy-2-butanone-4-phosphate (DHBP) with 5-amino-6-ribitylamino-2,4(1*H*,3*H*)-pyrimidinedione (also called 5-aminoribityluracil, 5-A-RU) first generates a Schiff base **1** (Figure 8A). Elimination of the phosphate group results in an enol that can tautomerize to a ketone, which can condense with the ribityl-amine. The release of water results in the formation of 6,7-dimethyl-8-ribityllumazine (ribityllumazine). Flavophos may act as a mimic of DHBP, potentially resulting in the formation of compounds **2**, **3**, or **4** (Figure 8B). Several observations support this mechanism of inhibition. When flavophos was added in increasing concentrations to a reaction containing DHBP, 5-A-RU, and *Ec*LS the production of ribityllumazine ceased (Figure S10) and two products were detected by LC-MS that are consistent with compounds **2** or **3** (both have m/z 437.1079) and **4** (m/z 419.0973) (Figures S11 and S12). Interactions between the phosphonyl moiety of flavophos and a conserved Arg residue in lumazine synthase (Arg127 in *Bsu*LS) may stabilize these intermediates and result in a bi-substrate analog inhibitor.

**Figure 8.**
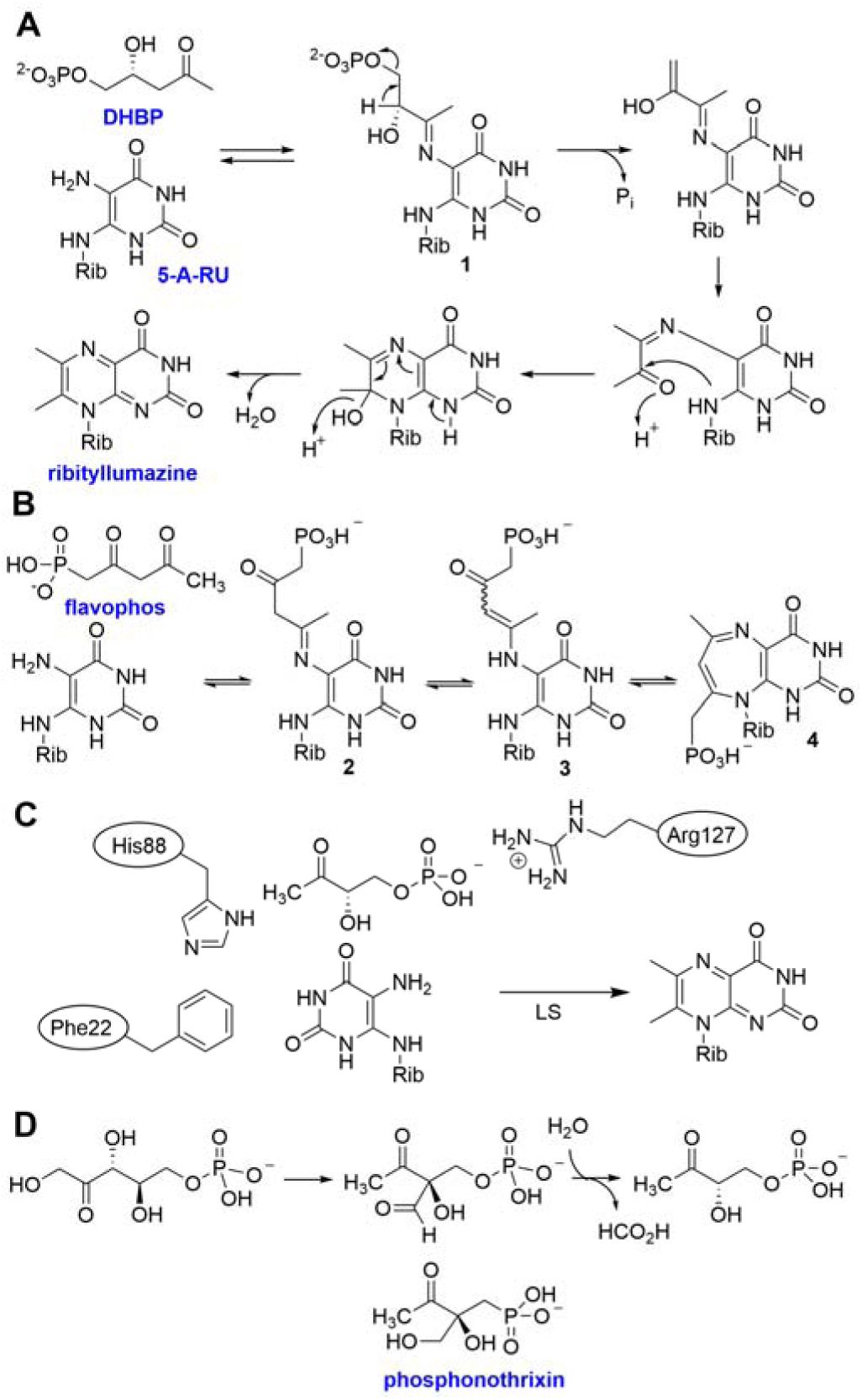
Mechanisms of LS catalysis and inhibition of LS by flavophos. **A**) DHBP and 5-A-RU are converted by lumazine synthase to ribityllumazine. **B**) The analogous reaction of flavophos with 5-A-RU may result in the formation of compounds **2**-**4**, resulting in LS inhibition. Masses consistent with compounds **2**-**4** were observed (Figures S11 and S12); **4** would be anti-aromatic if planar and might be expected to be unstable. Thus, a different tautomer may be favored or the compound attains a non-planar conformation as suggested by molecular mechanics energy minimization (Figure S13). **C**) Active site residues in *Bsu*LS implicated in binding substrates DHBP and 5-A-RU during catalysis.^49,63^ **D**) Reaction catalyzed by DHBP synthase and the structure of phosphonothrixin that inhibits the enzyme by mimicking the structure of an intermediate during catalysis.

## Discussion

Riboflavin (vitamin B2) is required by all forms of life and is the universal precursor of flavin mononucleotide (FMN) and flavin adenine dinucleotide (FAD). While in most eukaryotes riboflavin is acquired from the diet, in bacteria, archaea, and plants it is biosynthesized through a conserved biosynthetic pathway from guanidine triphosphate (GTP) and ribulose-5-phosphate.^66^ Medicinal chemistry groups have devised inhibitors of various riboflavin biosynthetic enzymes.^67–69^ Many of these inhibitors are substrate analogs that have been utilized for structural biology studies to understand the mechanistic details of catalysis.^70,71^ Surprisingly, to the best of our knowledge only one natural product has been reported that targets riboflavin biosynthesis, which coincidentally is another phosphonate, phosphonothrixin.^72^ This compound was recently shown to inhibit DHBP synthase by mimicking an intermediate that is formed during catalysis (Figure 8D).^73,74^ As such, phosphonothrixin has promising herbicidal activity.^72^ The structure of flavophos also resembles intermediates in the reaction mechanism proposed for LS,^39^ and while more detailed studies of the inhibition mechanism are warranted, the data in this study suggest that flavophos may form an adduct with the co-substrate 5-A-RU. The hypothesis that riboflavin biosynthesis is the target of flavophos is also supported by a gene encoding LS that is present in its BGC and by the observed protection to flavophos that is achieved by expression of LSs from various species. Our attempts to resolve whether resistance is conferred because of LS enzymatic activity or by sequestration of flavophos inside a protein cage were inconclusive. On the one hand *Mt*LS that does not form such a cage offered protection. On the other hand, so did mutants of *Bsu*LS that have very low activity. We note that no mutants of LS have been reported that completely abolished activity, consistent with the hypothesis that the enzyme acts mainly by controlling the orientation of the reactants but does not use side chains to actively catalyze the reaction.^63^

## Conclusion

Genome mining of phosphonate biosynthetic pathways that utilize 2-Pmm identified previously unknown phosphonate BGCs in *Burkholderia*. Reconstitution of the pathway in *E. coli* and subsequent *in vitro* studies resulted in discovery of a new compound with antibacterial activity. Structural and synthetic studies demonstrated the compound to be 2,4-dioxopentylphosphonic acid. Its antibacterial activity could be eliminated by over-expression of *bsfB*, a self-immunity gene within the BGC that encodes a lumazine synthase. The new phosphonate was shown to inhibit lumazine synthase in vitro, and overexpression of lumazine synthases from various organisms similarly conferred resistance. The phosphonate natural product was therefore termed flavophos. The biosynthetic pathway to flavophos involves an enzyme of the DUF849 family with sequence homology to β-keto acid cleavage enzymes. This enzyme (BsfD) catalyzes different chemistry than canonical members of this family, and co-crystal structures provide explanations for the observed activity. These findings will facilitate future efforts to elucidate pathways that involve homologs of BsfD.

## Experimental Procedures Methods and Materials

Type II restriction enzymes, Phusion® HF polymerase, Q5 polymerase, and the HiFi Assembly® kit were purchased from New England Biolabs (NEB). Primer DNA oligos were purchased from Integrated DNA Technologies (IDT), and synthetic gene fragments with codon optimized genes for expression in *E. coli* were purchased from Twist Bioscience (Table S4). For some experiments genomic DNA (gDNA) from *Burkholderia* sp. Bp9002 (Wagner and Sahl laboratories, University of Northern Arizona) was isolated with MoBio UltraClean Microbial DNA Isolation Kit and used for amplification of genes using the appropriate primers (Table S4). Plasmids were assembled using Gibson Assembly® (NEB) (Table S5).

LB medium components were purchased from Sigma Aldrich, Fisher Scientific, Alfa Aesar, and Acros Organics. Ampicillin (Amp), kanamycin (Kan), chloramphenicol (Cm), L-arabinose (ara), and isopropyl-β-D-thiogalactopyranoside (IPTG) were purchased from GoldBio and used at final concentrations of 100 μg/mL, 50 μg/mL, and 34 μg/mL, 0.2%, and 0.1 mM, respectively. Streptomycin (Sm) was purchased from Sigma Aldrich and used at concentrations of 100 μg/mL. Protein purification buffer components, HEPES, MOPS, glycerol, sodium chloride, imidazole, betaine, and D-sorbitol were obtained from Sigma Aldrich. FMN and FAD were purchased from Cayman Chemicals. Acetyl-CoA (lithium salt), and CoASH were purchased from CoALABio and used without further purification. 2-^13^C-acetic anhydride, lithium acetoacetate, and (-)-riboflavin were purchased from Sigma Aldrich. HisPur Ni-NTA resin was acquired from Thermo Fisher. PD-10 desalting columns were obtained from Cytiva. D_2_O was purchased from Cambridge Isotope Labs.

## Nuclear Magnetic Resonance spectroscopy

Spectra were recorded primarily on an Agilent NMR instrument operating at 599.70 MHz equipped with a OneNMR probe for ^1^H, ^13^C (150.82 MHz), and ^31^P (242.77 MHz) measurements. Data were acquired on a console operating VnmrJ 4.2. Where indicated, samples were acquired on either a Bruker NMR instrument operating at 600.13 MHz equipped with a BBO prodigy probe for ^1^H, ^13^C (150.92 MHz), and ^31^P (242.92 MHz) measurements, or a Bruker NMR instrument operating at 500.35 MHz equipped with a BBFO CryoProbe for ^1^H, ^13^C (125.83 MHz), and ^31^P (202.54 MHz) measurements. Data were acquired on a console operating Bruker TopSpin 4.1.0 or 3.6.2, respectively, and IconNMR for automated acquisitions using a SampleXpress autosampler. Data were acquired on a spectrometer with gradient/pulse-shaping capabilities and samples were held at 22-25 °C during acquisition without spinning. Samples were referenced to internal peaks (^1^H and ^13^C), or an external standard of 85% H_3_PO_4_ for ^31^P.

Quantitative NMR (qNMR) experiments were performed to determine flavophos concentration after chemical synthesis and purification. A reference standard of 1.75 mM maleic acid was prepared in D_2_O and the ^1^H NMR spectrum was recorded. The methylene protons at C1 (next to the phosphorus atom) were used to determine the concentration of flavophos solutions for the bioactivity assays. To determine the best peak of flavophos for quantification, all exchangeable protons in flavophos were replaced by deuterium using a solution in 100% D_2_O. The corresponding protons producing the doublet signal at C1 did not exchange after 24 h of D_2_O incubation whereas the protons at C3 exchanged.

## Construction of plasmids for co-expression and indicator strain testing

PCR products of *bsfD* homologs were inserted into the linearized vector of pETDuet_psfC generated by PCR, by Gibson Assembly (NEB). Gene products for *bsfE* and/or *bsfB* were ligated into linearized pCDFDuet vectors digested with *Nco*I/*Not*I for MCS1 insertion of *bsfE* homologs, or *Nde*I/*Pac*I for MCS2 insertion of *bsfB* homologs. For plasmids used in the indicator strain, pBAD-HisA was digested with *Nco*I/*Xho*I and ligated with compatible PCR products of *bsfB*, *Ec*LS (RibE; the naming of LS genes varies in different organisms), and *Bsu*LS (RibH), whereas *Mycobacterium tuberculosis* LS (*Mt*LS; RibH) was inserted in a pET29b plasmid. The *A. aeolicus* and *M. tuberculosis* enzymes were codon optimized for expression in *E. coli* and cloned into pET29b plasmids by Genscript for individual expression and purification of RibBA (3,4-dihydroxy-2-butananone-4-phosphate synthase/GTP cyclohydrolase II and lumazine synthase).

## Co-expression conditions and isolation of cell lysate and spent media

For preparation of cell lysate containing 3-OBPn, the resulting aqueous fraction after clarification was diluted with water to 80 mL total volume (arbitrary initial volume), prior to concentration via SpeedVac. Material was removed when 1 - 5 mL remained, to avoid decomposition of 3-OBPn into 2-OPP. The lysate was immediately frozen in liquid nitrogen and stored at -80 °C or used directly for NMR and experimental assays (*vide infra*).

## Generation of an SSN for PsfB

The sequence of PsfB from *Pseudomonas syringae* PB-5123 (AFM38993.1) was utilized as a BLAST search returning the top 1000 nearest hits. An arbitrary cutoff was chosen at 70% identity to generate an initial SSN. A full network was visualized and submitted through the GNN toolkit to generate a Pfam-associated network. Clusters within the SSN which contained a member of the PEP_mutase family were extracted. Sequences of PsfB proteins from these GCF members were then utilized as an additional BLAST query, and the resulting sequences were collated into a single list and duplicates removed. This new list of sequences was re-submitted through the SSN tools to generate another SSN. An alignment score of approximately 107 (50% identity) was chosen that segregated each family based on difference in gene cluster content.

Gene clusters from each family were analyzed by either the genome neighborhood diagram tool (GNN) or RODEO if the sequences were absent from UniProt.

## Bioactivity assays against *E. coli* WM6242 **λ**(DE3)

For bioactivity testing against *E. coli* WM6242 λ(DE3), agar plates were prepared by autoclaving LB agar (at 1.5% agarose) and subsequently cooling to 50 °C. To this cooled agar was added IPTG to 1 mM, and approximately 25-30 mL of agar was poured into sterile Petri dishes. Plates were stored at 4 °C for less than 6 months. Meanwhile, 6 mm paper filter disks (BD) were arranged on cooled agar plates and impregnated with 5-10 μL of material. A starter culture of *E. coli* WM6242 λ(DE3) was seeded from a single colony and grown at 37 °C until slightly turbid (OD_600_ 0.4-0.6). This culture was used to inoculate top agar consisting of molten LB agar (at 0.7% agarose) in a 1:100 dilution immediately prior to pouring on top of the filter disks. Approximately 6-8 mL of top agar was utilized per plate to generate a thin film of inoculated agar.

## Purification of BsfD

*Bs*BsfD and *Bm*BsfD were expressed and purified in a manner similar to that described for members of the BKACE protein family.^36^ *E. coli* BL21(DE3) harboring pET28a plasmids with either codon optimized gene was grown in LB media containing 50 μg/mL kanamycin overnight. Expression cultures were prepared containing 1 L TB media made up of 20 g tryptone, 24 g yeast extract, 0.4% glycerol containing 0.5 M sorbitol and 5 mM betaine, added prior to autoclaving, diluted with water to 900 mL. After cooling, 100 mL of 10 × phosphate buffer (0.017 M KH_2_PO_4_, 0.072 M K_2_HPO_4_, final concentration) was added, followed by addition of kanamycin to 200 μg/mL. To 1 L of expression culture was added 10 mL of overnight cultures and incubated at 37 °C until an OD of 0.8, at which point IPTG was added to 0.5 mM. After induction the cultures were cooled to 18 °C and continued to incubate with shaking overnight.

Cells were collected by centrifugation and yielded 30 g/L for *E. coli* BL21(DE3) containing pET28a_His-*Bm*BsfD and 90 g/L for cells containing pET28a_His-*Bs*BsfD. Because of the higher cell yield for the *Bs*BsfD culture, all in vitro experiments and structural characterization were performed with this ortholog. Cell pellets were resuspended in lysis buffer containing 50 mM Na_2_HPO_4_, pH 7.5, 10 mM imidazole, and 300 mM NaCl; DNase and lysozyme were added to 1000 U and 3.5 mg/mL buffer, respectively, followed by incubation at 4 °C for 30 m on a rocker. Cells were ruptured by passage through a chilled French pressure cell (Thermo Electron Corporation) twice, and cell debris was removed by centrifugation. Clarified supernatant was applied to HisPur resin pre-equilibrated with lysis buffer at 4 °C. The resin was washed twice with lysis buffer, followed by the same buffer containing 20 mM imidazole. Protein was eluted with buffer containing 250 mM imidazole and 10% glycerol. Fractions containing protein were pooled, concentrated, and applied to a PD-10 desalting column (Cytiva) equilibrated with storage buffer (50 mM Na_2_HPO_4_, 500 mM NaCl, and 10% glycerol at pH 7.5). Protein was eluted and stored at −80 °C.

## Enzyme assays of BsfD utilizing ^13^C-AcCoA

Enzyme assays were performed to determine BsfD activity with (2-^13^C)-AcCoA to evaluate methyl group transfer to acetoacetate. Utilizing cell lysate derived from co-expression of *psfABC* as a source of 3-OBPn, the lysate was neutralized with addition of phosphate buffer to 40 mM, pH 7.4, then (2-^13^C)-AcCoA was added to 10 mM. The (2-^13^C)-AcCoA solution (stored at 140 mM) is acidic, therefore the solution was neutralized with 1 M KOH to pH 7.4 prior to addition of enzyme. The reaction was initiated by addition of purified *Bs*BsfD to 90 μM. Reactions were set up in parallel with storage buffer *in lieu* of *Bs*BsfD, or lysate of cells expressing *psfAB in lieu* of *psfABC* (2-Pmm containing lysate, no 3-OBPn present).

After overnight incubation, samples were analyzed by ^31^P and ^13^C NMR spectroscopy. The sample containing lysate of cells expressing *psfABC* and *Bs*BsfD protein was spiked with 10 mg lithium acetoacetate (185 mM, final concentration). After additional ^13^C NMR acquisitions, a drop of acetone was added to the sample for additional data acquisition.

## Synthesis of (2,4-dioxopentyl)-phosphonic acid

Flavophos was prepared from diethyl-2-oxopropylphosphonate using a reaction described previously.^43^ After workup, the product was lyophilized and used directly for NMR spectroscopy and bioactivity experiments. The indicated couplings were observed and assisted in structural assignments. δ_P_ (242.78 MHz, D_2_O): 9.35 (s). δ_H_ (599.70 MHz, D_2_O): 2.40 (3 H, s, C(O)-C**H_3_**), 3.11 (2 H, d, ^2^*J*_HP_ 19.8 Hz, P-C**H_2_**-C(O)), 4.12 (2 H, s, C(O)-C**H_2_**-C(O)). δ_C_ (150.81 MHz, D_2_O): 27.32 (s, C(O)-**C**H_3_), 203.36 (CH_2_-**C**(O)-CH_2_), 206.76 (CH_2_-**C**(O)-CH_3_).

## Crystallization and structure determination of *Bs*BsfD with various ligands

Recombinant *Bs*BsfD was purified as described above but with the inclusion of an additional chromatographic step prior to crystallization. Briefly, following elution from nickel-affinity chromatographic resin, aliquots of purified *Bs*BsfD was treated with thrombin protease (Cytiva, USA) to remove the His_6_ affinity tag. After excision of the tag was complete, as determined by denaturing SDS-PAGE analysis, *Bs*BsfD was further purified by size-exclusion on a Superdex 75 column preequilibrated with buffer composed of 150 mM KCl and HEPES pH=7.5. Samples with the highest purity were pooled, concentrated to ∼30 mg/mL and subject to crystallization trials using an Art Robbins Gryphon liquid dispenser.

Crystals of ligand-free *Bs*BsfD were formed using 1.6 M lithium sulfate, Tris-HCl, pH=8.5 and grew over the course of 2 days. Co-crystals of *Bs*BsfD incubated with CoA grew under the same conditions. Crystals were soaked in a solution of the crystallization medium supplemented with 30% glycerol (v/v) prior to vitrification by direct immersion into liquid nitrogen. For co-crystallization with substrate, enzymatically generated 3-OBPn (and the decarboxylated product) was incubated with *Bs*BsfD and acetyl-CoA and subject to co-crystallization efforts. Crystals were formed using 100 mM calcium acetate, 100 mM sodium acetate, pH=4.5 and varying concentrations of polyethylene glycol 4000. Diffraction quality crystals were subsequently incubated in the crystallization media supplemented with additional aliquots of 3-OBPn for short (6 h) or long (48 h) time increments before vitrification into liquid nitrogen for data collection. All diffraction data were collected at the Life Sciences Collaborative Access Team (LS-CAT) at Argonne National Labs (beamline 21-G, wavelength = 0.97857 Å). All diffraction data were index, scaled, and integrated using either XDS^75^ or autoPROC.^76^ Crystallographic phases were determined by molecular replacement as implemented in PHASER^77^ using the atomic coordinates of *Ra*Kce (PDB code 3C6C) as a search probe. The initial molecular model was manually rebuilt in COOT^78^ using the sequence of *Bs*BsfD prior to iterative rounds of refinement using REFMAC5^79^ interspersed with further manual building until convergence as determined by the free R factor and Ramachandran analysis. Final crystallographic statistics are listed in Table S2.

## Resistance mutant generation

An overnight culture of the phosphonate indicator strain *E. coli* WM6242 λ(DE3) was inoculated in 2 mL of LB containing 32 µM flavophos. After overnight incubation at 37 °C, the culture was cloudy indicating resistant bacteria growth. This batch of liquid culture was streaked on an LB-agar plate containing 32 µM flavophos followed by overnight incubation. The next day, colonies were grown in fresh LB solution (2 mL) containing 32 µM flavophos. After overnight growth, a glycerol stock was made to preserve the resistant bacteria (sample A). Another set of spontaneous resistance mutants was obtained by using a culture grown overnight at 32 µM flavophos, dilution 100-fold and inoculation of a 2 mL LB culture containing 1 mM flavophos. After 24 h, the resistant culture was stored for future use (sample B). A final spontaneous resistant mutant was obtained by picking a single colony that appeared in the zone of growth inhibition when *E. coli* WM6242 λ(DE3) was plated on LB agar containing 1 mM flavophos. A glycerol stock was then prepared the next day and stored for future use (sample C). Each glycerol stock containing the three different flavophos-resistant cultures was streaked on to an LB-agar plate containing 0.1 mM flavophos and 1 mM IPTG and incubated overnight at 37 °C. Next day, four single colonies from each sample were subjected to colony PCR using primers that amplify a region of the genome that encompasses the phosphonate uptake genes (*phnCDE*) and the lumazine synthase gene (*ribE*). Each PCR amplified product was then submitted for sequencing. Additionally, another four single colonies from sample B sample grown were grown at 0.7 mM flavophos and the genomic DNA was extracted. Each gDNA sample was then submitted for full genome sequencing.

## In vitro inhibition of *Ec* lumazine synthase, RibE

N-terminal His-tagged *Ec*RibE and *Ec*RibB were purified using Ni-affinity chromatography. Briefly, *E. coli* BL21(D3) harboring the plasmid pRSFDuet-His6_EcRibE or pRSFDuet-His6_EcRibB was grown at 37 °C until OD600 reached 0.6-0.8. IPTG (0.4 mM) was used to induce the expression of RibE or RibB and the culture was further grown at 18 °C overnight. Cells were harvested at 5,000 ×g at 4 °C and resuspended in 50 mM HEPES, 300 mM NaCl, pH 7.6 containing 1 mM TCEP and 1 mM PMSF. The solution was then sonicated for 3 min with a pulse cycle of 2 s on and 5 s off at 50% amplitude. The lysate was then centrifuged at 60,000 ×g for 45 min and the clear supernatant was passed through a Ni-affinity column. The resin was washed twice with 25 mM imidazole, 300 mM NaCl, pH 7.6, and 10% glycerol and the protein was eluted with 200 mM imidazole, 300 mM NaCl, pH 7.6, and 10% glycerol. The buffer of the eluted protein was then exchanged three times with 50 mM HEPES, 300 mM NaCl, pH 7.6 using a 10K Da cutoff Amicon.

In vitro reactions to assay LS activity were performed in a 200-µL tube containing 0.15 mM 5-A-RU, 0.1 mM ribulose-5-phosphate, 20 mM MgCl_2_, 5 mM DTT, 11 µM RibB, and 50 mM Tris-HCl pH 7.6. After incubation for 30 min at 37 °C, EDTA was added (10 mM final concentration), followed by variable concentrations of flavophos (0, 0.042, 0.125, 0.375, 0.6, 1, and 3 mM) and 6 µM RibE (*Ec*LS) to start the reaction. Aliquots of 25 µL were taken at 15 min, 30 min, and 60 min and quenched with 6 µL of 20% TCA to precipitate proteins. The solutions were then centrifuged, and the supernatant analyzed by LC-MS analysis using a poroshell EC-C18 column with the analytes measured in negative mode.

## Resistance conferred to *E. coli* by *Mt* lumazine synthase

An overnight culture of the phosphonate sensitive strain WM6242 λ(DE3) expressing *Mt*LS was used to inoculate (1:100 ratio) 10 mL top LB-agar (1% agar) containing 50 µg/mL kanamycin and 0.1 mM IPTG and poured over 4 mL of set LB-agar (1.5% agar) also containing 50 µg/mL kanamycin and 0.1 mM IPTG. Two separate 200 µL tubes were placed in the top agar to create two wells. The agar was allowed to set for 30 min and 5 µL solutions of 100 µM and 200 µM flavophos were added to each well. The plates were covered and incubated at room temperature overnight upon which a lawn of bacterial growth were observed without any visible inhibition zones. As a control, the same protocol was followed using only the indicator strain and four flavophos concentrations including 100 µM showing clear inhibition zones at 25 and 100 µM.

## Generation of LS and BsfD mutants

The backbone of pBADCDF was amplified by PCR, and the desired variant of lumazine synthase was ordered as a G-block from Twist (Table S4) and inserted into the plasmids by Gibson Assembly (NEB). For the pBADCDF_AaLs plasmid, pBADCDF was digested with *Nco*I/*Pst*I and ligated with compatible PCR products of AaLs. Similarly, the BsfD-F166Y mutant was made from a G-block and inserted into pET15b.

## Supporting Information

Experimental procedures, Figures S1-S13 showing mass spectrometry, bioactivity, and NMR data, and Tables S1-S5 with gene sequences, crystallographic information, gene sequence, primers, and plasmids used.

## Notes

The authors declare no competing financial interests.

## Supporting information

Cytoscape files

## Acknowledgments

The authors thank David M. Wagner and Jason W. Sahl at the University of Northern Arizona for providing *Burkholderia* strains that have the *bsf* BGC in their genomes.

