## Supporting Information for "Discovery of the Phosphonate Flavophos Produced by *Burkholderia*"

**Table of Contents** **Page #**

**Figure S1.** General activity observed for members of the BKACE family of proteins. **S2**
**Figure S2.** Assays of BsfD using 2-13C-AcCoA analyzed by 31P NMR spectroscopy. **S3**
**Figure S3.** $^{1}$H-$^{31}$P HMBC NMR of enzymatically produced and synthetic flavophos. **S4**
**Figure S4.** Structure-based multiple sequence alignment of *Bs*BsfD. **S5**
**Figure S5.** Anomalous difference Fourier maps near the Zn absorption edge. **S6**
**Figure S6.** LigPlot of the active site of *Bs*BsfD in the vicinity of the active site metal. **S7**
**Figure S7.** LigPlot of the active site of *Bs*BsfD in the vicinity of the CoA. **S8**
**Figure S8.** Bioactivity testing of flavophos *via* agar diffusion assays. **S9**
**Figure S9.** Bioactivity testing of flavophos against LS mutants *via* agar diffusion assays. **S10**
**Figure S10.** LC-MS traces of the in vitro inhibition assays of *Ec*LS **S11**
**Figure S11.** Observed LC MS/MS fragmentation consistent with adduct 2 or 3. **S12**
**Figure S12.** Observed LC MS/MS fragmentation consistent with putative adduct 4 **S13**
**Figure S13.** Minimized geometry for potential adduct 4 **S14**
**Table S1.** Gene names and accession numbers in this study **S15**
**Table S2.** Crystallography data collection and refinement statistics. **S16**
**Table S3.** Summary of mutations in *E. coli* WM6242 against flavophos **S17**
**Table S4.** Primers for plasmid assembly and codon-optimized genes **S19**
**Table S5.** Plasmids and microorganisms used in this study. **S24**
**References** **S26**

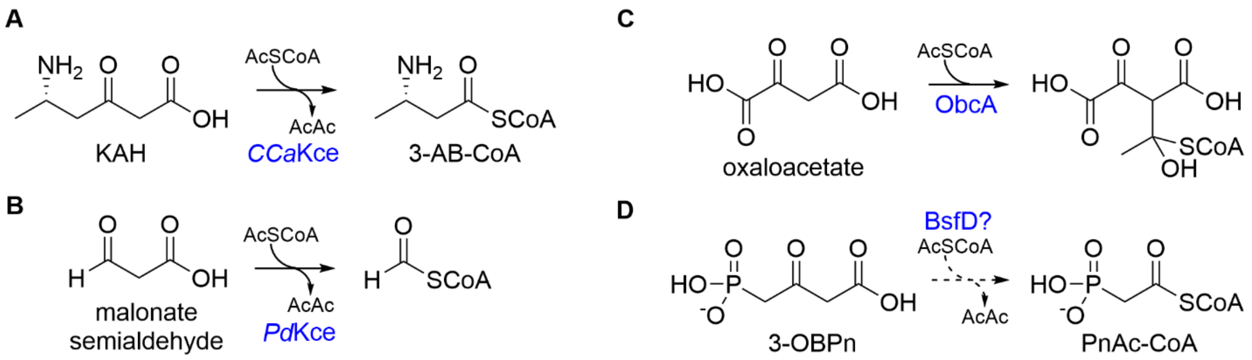


**Figure S1.** General activity observed for members of the BKACE family of proteins. **A**) The reaction catalyzed by *CCa*Kce as part of lysine fermentation converting 3-keto-5-aminohexanoic acid (KAH) and acetyl-CoA to acetoacetate (AcAc) and 3-aminobutyryl-CoA (3-AB-CoA).[1] **B**) Reaction catalyzed by *Pd*Kce.[2] The reaction shown was used for a synthetic biology pathway; the physiological reaction catalyzed by *Pd*Kce is not known. **C**) Reaction catalyzed by ObcA.[3] This enzyme is related to BKACEs and catalyzes only the first step of canonical BKACE chemistry, the addition of the enolate of oxaloacetate to acetyl-CoA. In canonical BKACE chemistry, the CoASH is expelled next resulting first in a Claisen condensation product, which is followed by addition of the CoASH to the ketone in the original β-keto acid to initiate a retro-Claisen reaction (see Figure 7A of the main text). **D**) Reaction predicted to be catalyzed by BsfD if the chemistry were analogous to *CCa*Kce or *Pd*Kce. 3-OBPn and AcCoA would be transformed into phosphonoacetyl-CoA (PnAc-CoA).

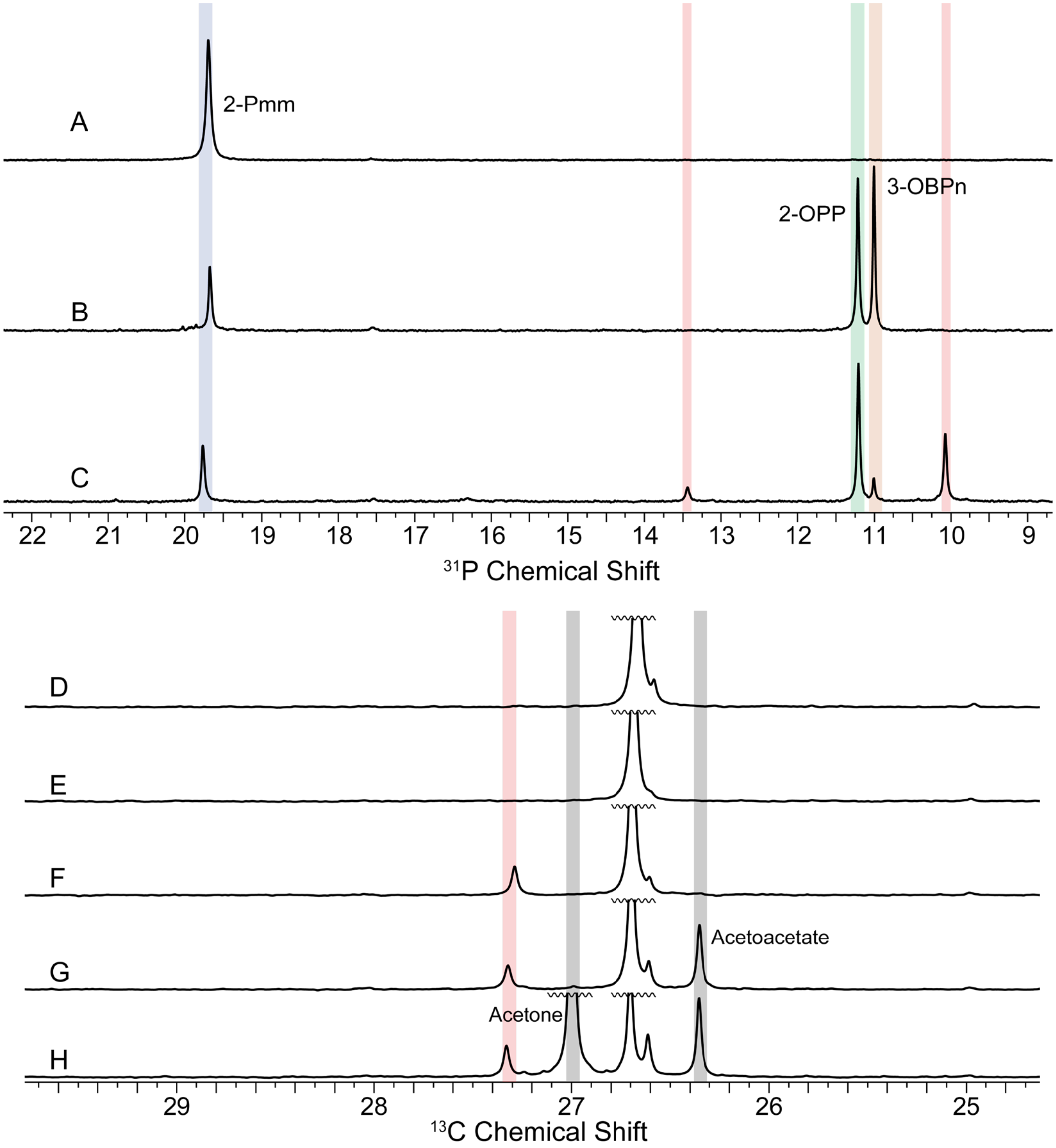


**Figure S2.** Assays of BsfD using 2-$^{13}$C-AcCoA analyzed by $^{31}$P NMR spectroscopy. 2-$^{13}$C-AcCoA was added to **A**) lysate from *E. coli* cells co-expressing *psfAB* and purified *Bs*BsfD, **B**) lysate from cells co-expressing *psfABC*, and **C**) lysate from cells co-expressing *psfABC* with added purified *Bs*BsfD. New peaks are indicated in red. **D-F**) $^{13}$C NMR spectroscopy analysis of samples **A-C**, respectively, shows a new peak at ~27.4 ppm (red) that is only present in the sample containing lysate of cells expressing *psfABC*, incubated with 2-$^{13}$C-AcCoA and purified BsfD. **G**) Addition of acetoacetate to this sample shows a new peak at ~26.35 ppm. **H**) Addition of acetone shows a new peak at 27 ppm. The peak highlighted in red (the flavophos product) is not associated with acetoacetate or its degradation product acetone.

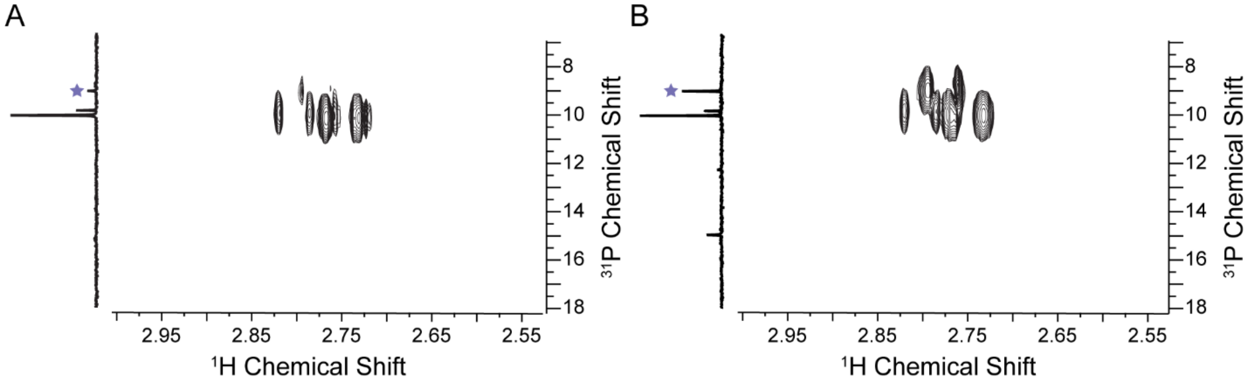


**Figure S3.** $^{1}$H-$^{31}$P HMBC NMR analysis of enzymatically produced and synthetic flavophos. **A**) $^{1}$H-$^{31}$P HMBC NMR spectrum of the sample in panel **C** of Figure 5 of the main text, and **B**) spiked with synthetic 2,4-dioxopentylphosphonic acid. A corresponding increase in intensity for the doublets corresponding with flavophos confirms they are derived from the same product.

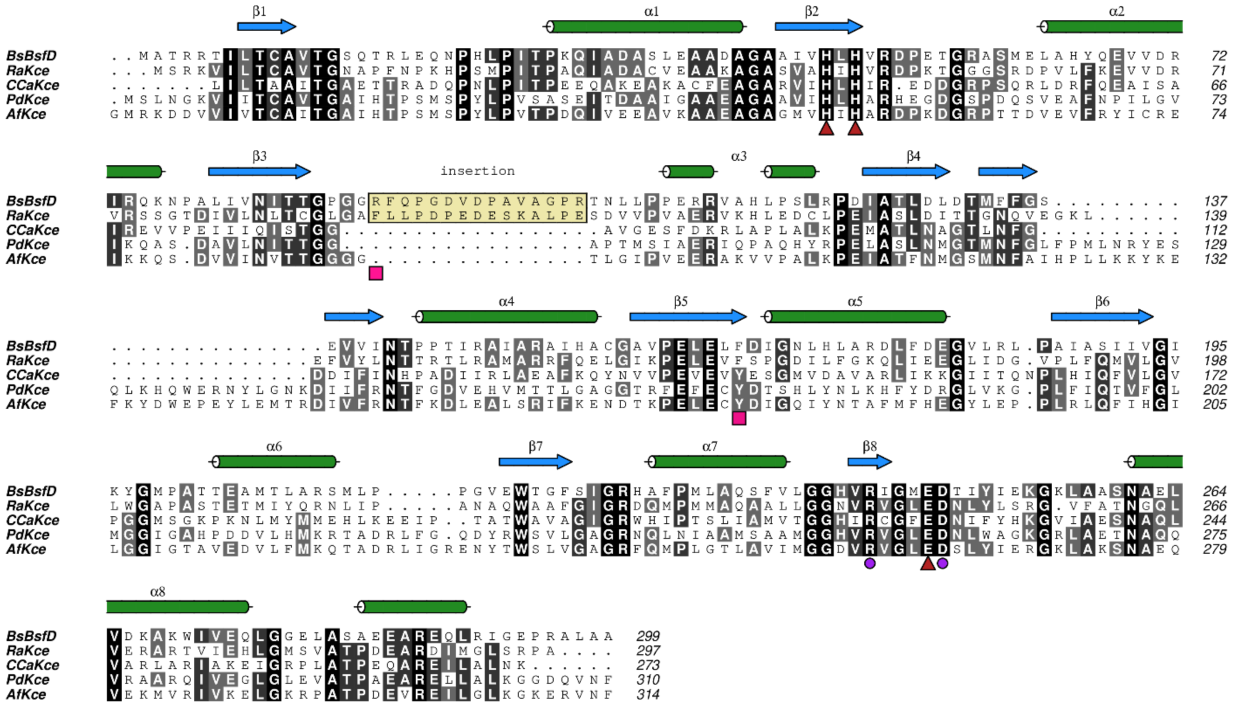

**Figure S4.** Structure-based multiple sequence alignment of *Bs*BsfD along with homologous enzymes that fall into the DUF849 superfamily and that have been functionally characterized. Residues involved in coordinating the divalent metal are labelled with a red triangle, conserved residues thought to be involved in acid-base catalysis in BKACE chemistry are marked with a purple circle. The Arg residue unique to *Bs*BsfD is marked with a pink square, as is the Phe that is a Tyr implicated in catalysis in several other canonical BKACEs (see main text).

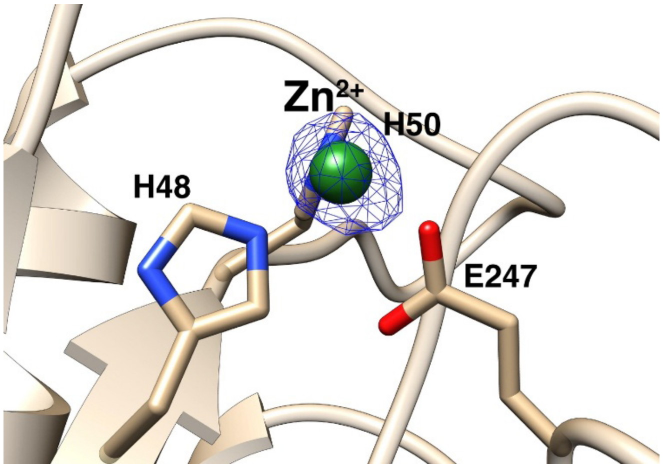


**Figure S5.** Anomalous difference Fourier maps calculated using data collected near the Zn absorption edge (1.1271 Å) and phased from the final model. The map is contoured at 8σ (blue mesh). The coordinates of the final structure are superimposed.

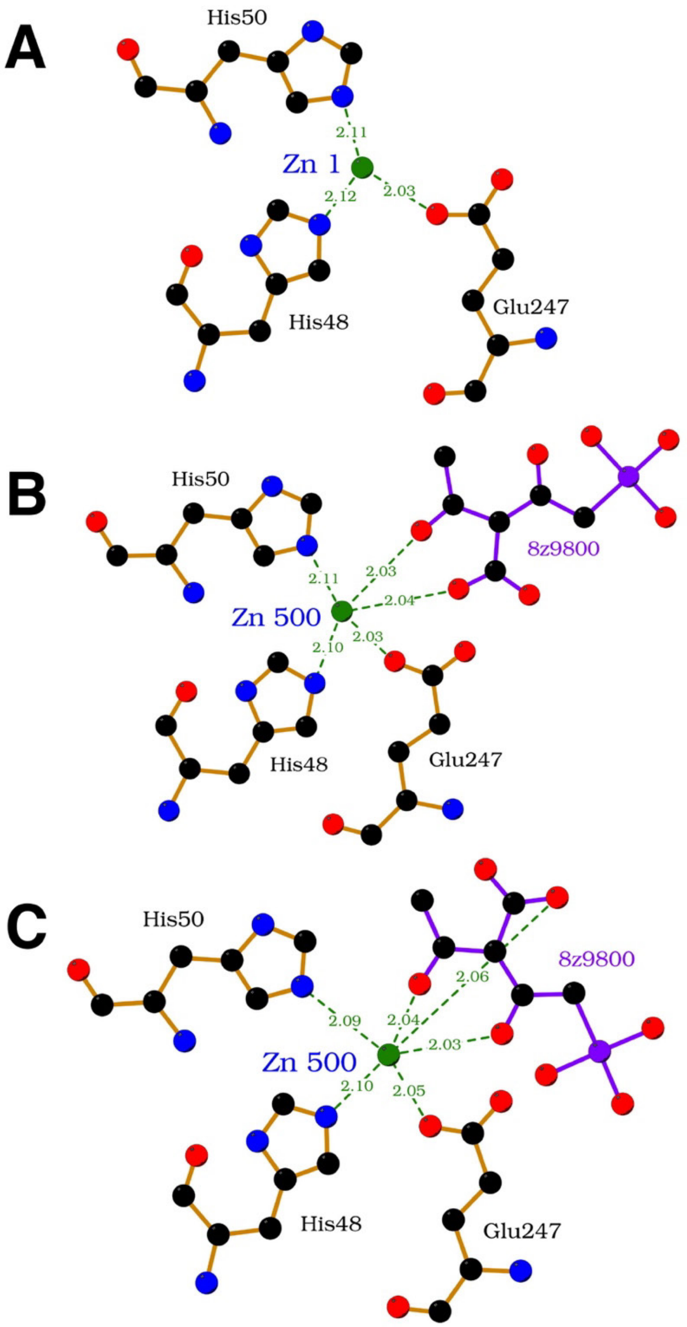


**Figure S6.** LigPlot of the active site of *Bs*BsfD in the vicinity of the active site metal derived from the coordinates of (A) the enzyme in the absence of ligands, (B) the enzyme co-crystallized with Ac-CoA and 3OBPn for 6 h, and (C) the enzyme co-crystallized with Ac-CoA and 3OBPn after 48 h. For clarity, the CoA is not shown in this Figure. A LigPlot focusing on the position of CoA is presented in Figure S7.

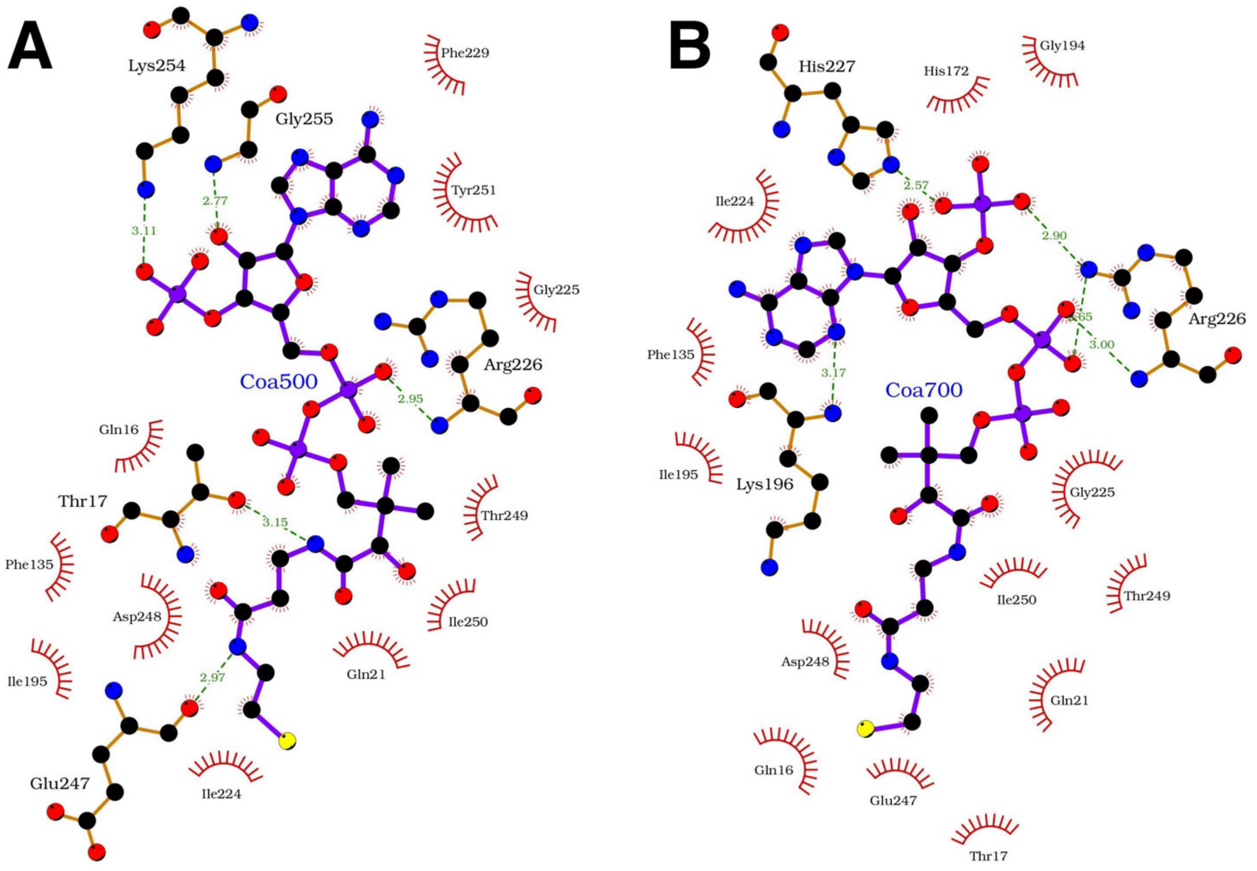


**Figure S7.** LigPlot figure of the active site of *Bs*BsfD in the vicinity of CoA derived from the coordinates of (A) the enzyme in the presence of CoA only, (B) the enzyme co-crystallized with Ac-CoA and 3OBPn for 48 h.

**A**

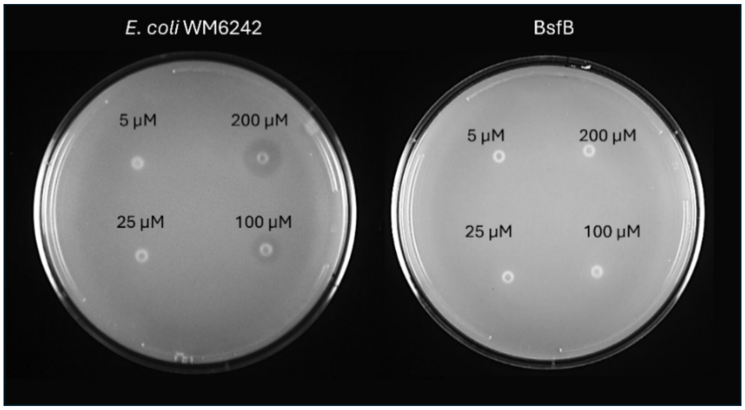


**B**

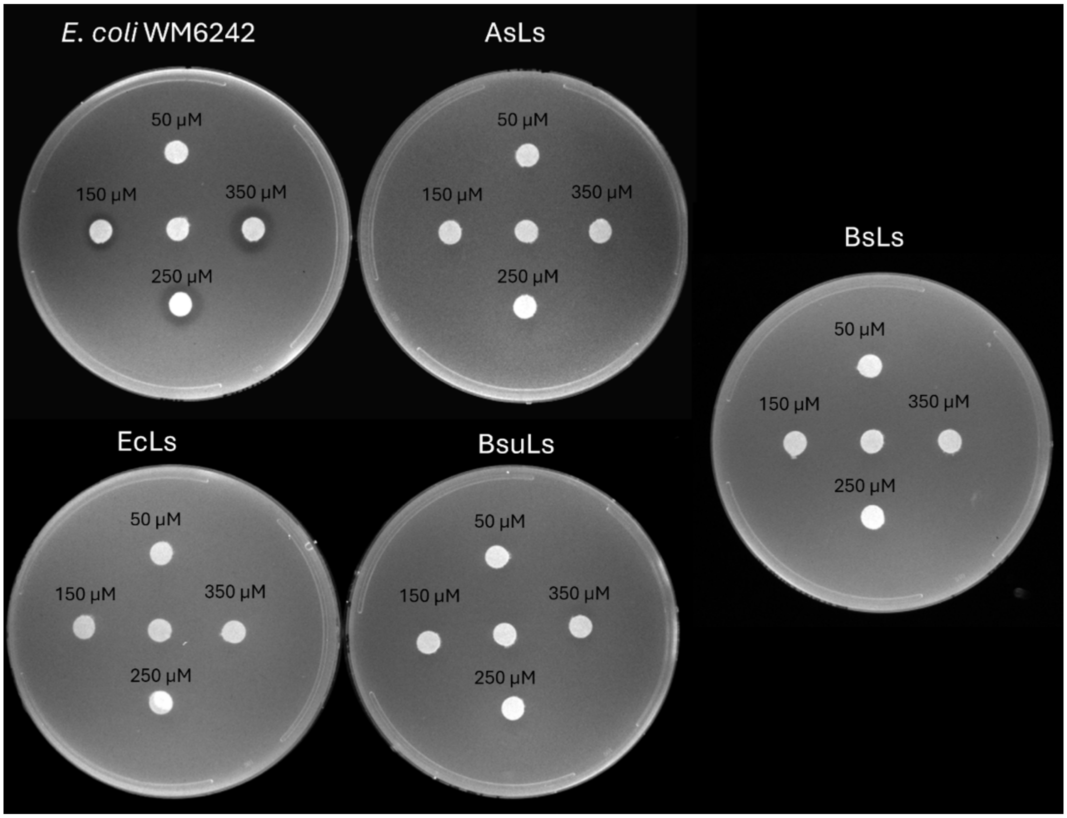


**Figure S8.** Bioactivity testing of flavophos *via* agar diffusion assays. **A**) Agar diffusion bioactivity assay of phosphonate sensitive strain WM6242 λ(DE3) (left) in the presence of IPTG and the strain transformed with pBAD-*Bs*BfsB (right) in the presence of arabinose (0.2%) and IPTG (1 mM). For each spot, 3 µL of a solution of flavophos with the concentrations shown per well. **B**) Various lumazine synthases (LSs) provide protection against flavophos when expressed in *E. coli* WM6242 λ(DE3). The images show the results of expression of the housekeeping LS of *Burkholderia* sp. BDU6 (*Bs*Ls), LS from *E. coli* (*Ec*LS), LS from *B. subtilis* (*Bsu*LS), and LS from *Aquifex aeolicus* (*Aa*LS) Every image shows plates where 2 µL of different solutions with the indicated concentrations of flavophos were placed on paper disks.

**A**

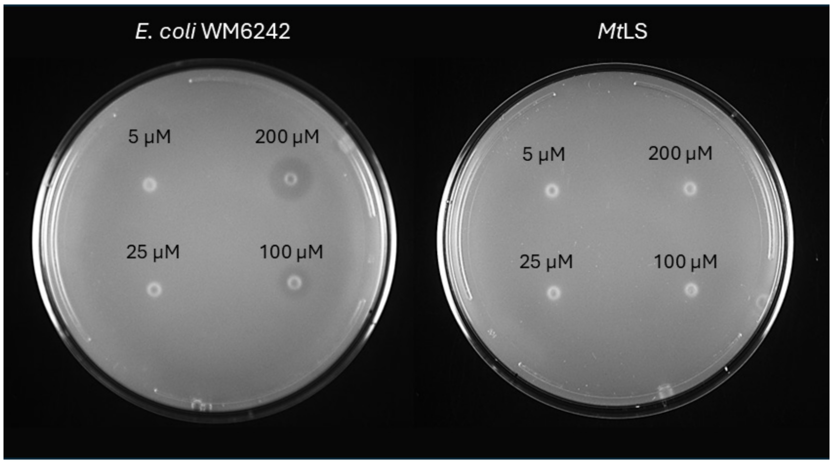


**B**

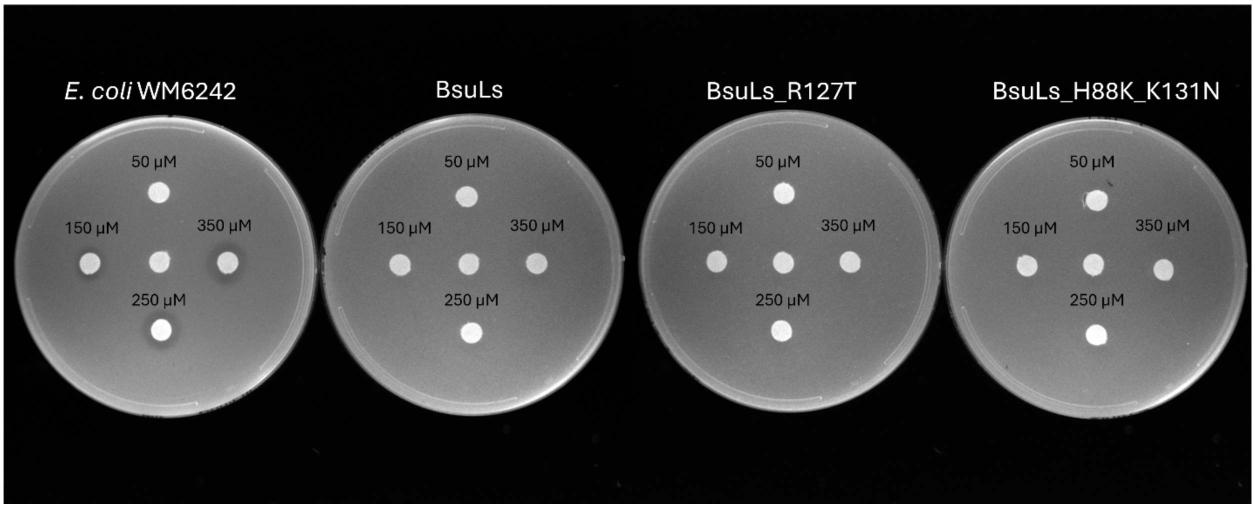


**Figure S9.** Bioactivity testing of flavophos against LS mutants *via* agar diffusion assays**. A)** Agar diffusion bioactivity assay of phosphonate sensitive strain WM6242 λ(DE3) (left) transformed with pET29b-*Mt*LS in the presence (right) of IPTG. For each spot, 3 µL of a solution of flavophos with the concentrations shown per well. **B)** Two lumazine synthase mutants from *B. subtilis* that have the lowest reported enzymatic activities[4] provide protection against flavophos when expressed in *E. coli* WM6242 λ(DE3). For each spot, 2 µL of a solution of flavophos with the concentrations shown was placed on the paper filter disks.

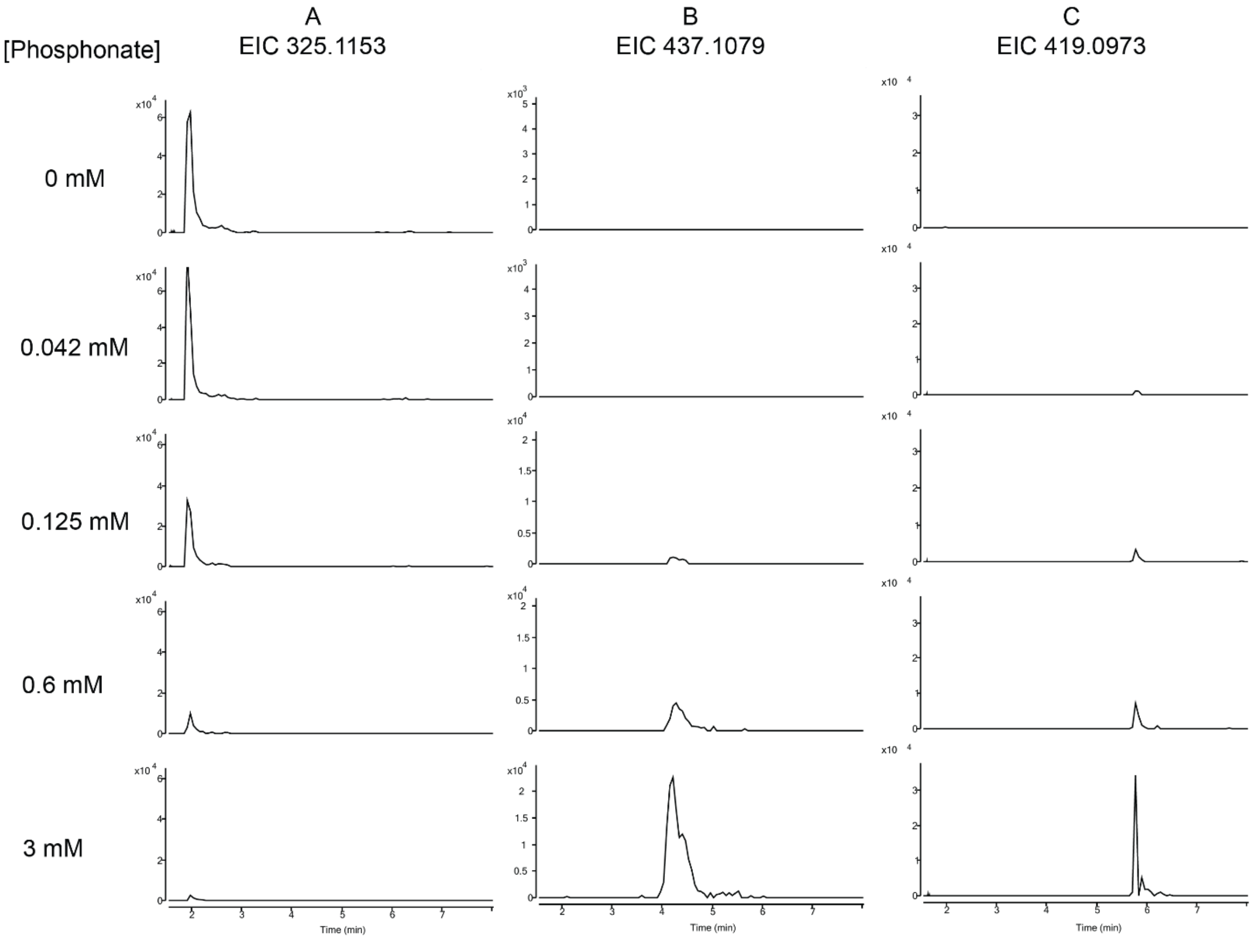


**Figure S10.** LC-MS traces of the in vitro inhibition assays of *Ec*LS showing the extracted ion chromatograms (EIC) for the $[M-H]^-$ ions corresponding to the formation of ribityllumazine (A), adduct **2** (B), and adduct **4** (C) with respect to increasing concentrations of flavophos. At 600 μM flavophos, ribityllumazine formation is nearly abolished and ions consistent of adducts **2** and **4** are observed. See Figures S11 and S12 for the mass spectra as well as fragmentation data.

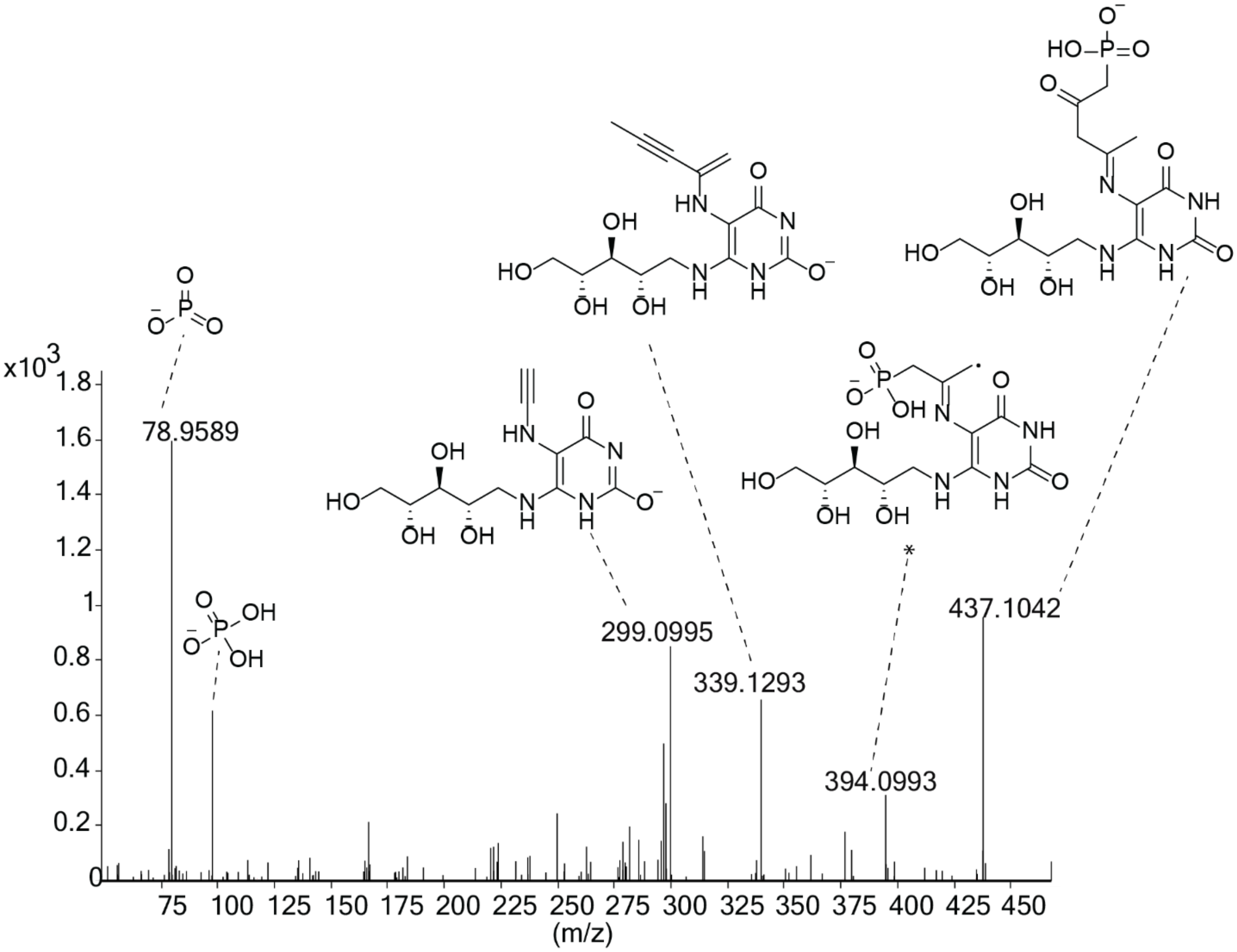


**Figure S11.** Observed LC MS/MS fragmentation consistent with adduct **2** or **3**. Some of the ions could be accounted for by more than one structure, e.g. the ion at 437 Da could come from the imine shown, the imine at C2 of flavophos, or their enamine analogs.

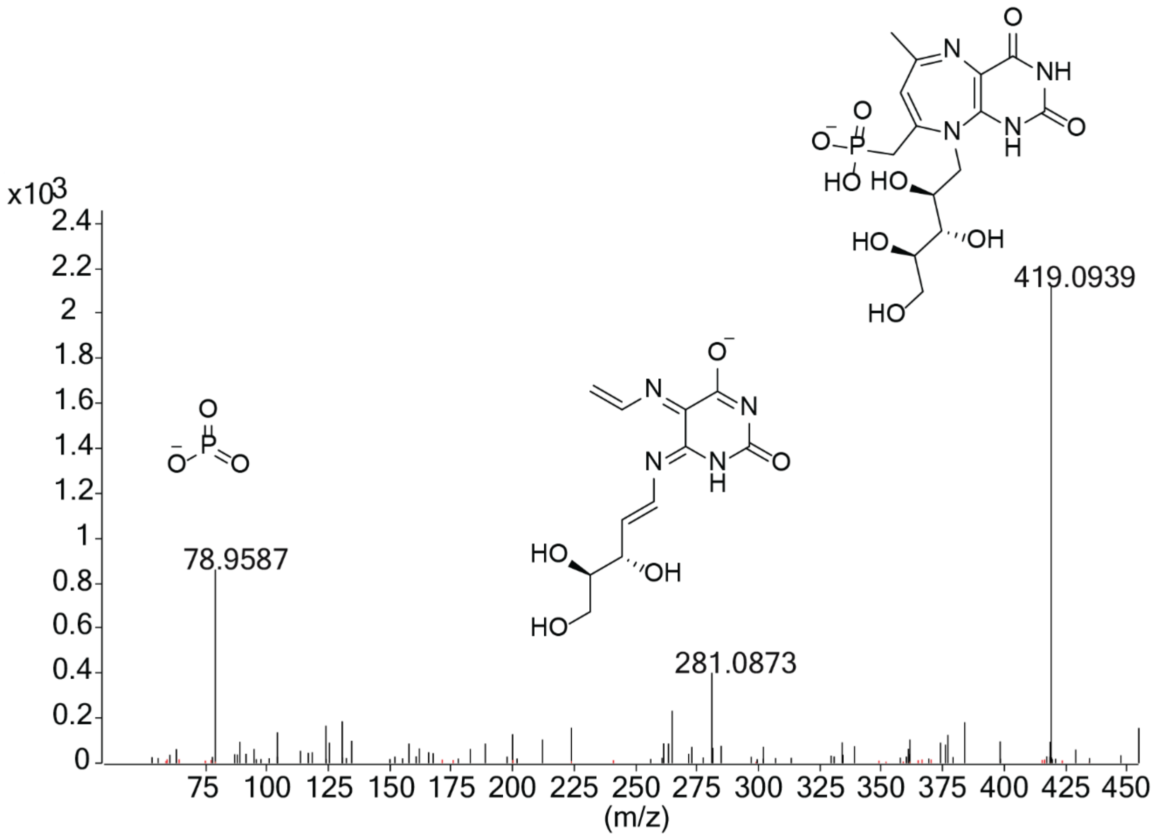


**Figure S12.** Observed LC MS/MS fragmentation consistent with putative adduct **4**.

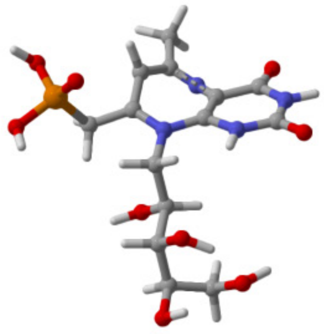

**Figure S13.** Minimized geometry for the potential adduct **4** in Fig. 7 showing that the compound would attain a non-planar geometry. The energy minimization calculation was performed using the free software Avogadro2[5] and the output was file was submitted to CYLview[7] for visualization.

**Table S1.** Gene names, accession numbers, and closest predicted function by UniProtKB/SwissProt or NCBI nonredundant database.

| Gene Name | Accession Number | Predicted Function (closest UniProtKB/SwissProt Hit) | % seqID to Closest Hit | Observed/Assumed Activity |
|---|---|---|---|---|
| *bsfA* | AOJ04873.1 | Fosfomycin PEP mutase (*Streptomyces wedmorensis*) | 57% | PEP mutase |
| *bsfB* | AOJ05795.1 | 6,7-Dimethyl-8-ribityllumazine synthase (*Metallosphaera sedula* DSM 5348) | 44% | -- |
| *bsfC* | AOJ04874.1 | PsfC (*Pseudomonas syringae* PB-5123) | 37.7 | PsfC-like |
| *bsfD* | AOJ04875.1 | 3-Keto-5-aminohexanoate cleavage enzyme (*Fusobacterium nucleatum*) | 35% | -- |
| *bsfE* | AOJ04876.1 | Drug/metabolite transporter (*Burkholderia* sp.) | 58% | Transporter |
| *bsfF* | AOJ04877.1 | 2-Phosphonomethylmalate synthase (*Streptomyces rubellomurinus*) | 64% | Pmm synthase |

**Table S2.** Data Collection and Refinement Statistics

| | ***Bs*BsfD (apo)** | **CoA** | **Ac-CoA+3OBPn (6h)** | **Ac-CoA+3OBPn (48h)** |
|---|---|---|---|---|
| **Data collection** | | | | |
| Space Group | I222 | I222 | $P2_12_12$ | $P2_12_12$ |
| Cell: a, b, c (Å) | 53.4, 120.2, 121.4 | 53.4, 119.6, 121.6 | 86.8, 109.6, 72.4 | 86.1, 110.5, 72.7 |
| Resolution (Å)[1] | 85.5 – 1.72 (1.75 – 1.72) | 48.9 – 1.49 (1.54 – 1.49) | 34.4 – 1.81 (1.84 – 1.81) | 37.0 – 1.74 (1.77 – 1.74) |
| Total reflections | 594,150 | 592,774 | 940,429 | 1,047132 |
| Unique reflections | 42,079 | 62,636 | 64,356 | 72,100 |
| $R_{sym}$ (%) | 7.2 (103.8) | 9.6 (57.7) | 9.9 (124.2) | 6.8 (121.2) |
| $I/\sigma(I)$ | 23.3 (2.1) | 13.4 (2.0) | 20.1 (2.3) | 27.6 (2.2) |
| Completeness (%) | 99.5 (97.8) | 99.8 (97.5) | 100 (100) | 100 (100) |
| Redundancy | 14.1 (10.8) | 9.5 (3.1) | 14.6 (14.5) | 14.5 (13.0) |
| CC (1/2) | 0.998 (0.763) | 0.997 (0.599) | 0.999 (0.779) | 0.999 (0.829) |
| **Refinement** | | | | |
| Resolution (Å) | 25.0 – 1.72 | 25.0 – 1.5 | 25.0 – 1.81 | 25.0 – 1.74 |
| No. reflections | 39,765 | 59,394 | 60,654 | 68,152 |
| $R_{work}$ / $R_{free}$[2] | 18.1 / 19.9 | 16.9 / 18.6 | 18.3 / 20.5 | 18.9 / 20.4 |
| Number of atoms | | | | |
| Protein | 2231 | 2243 | 4355 | 4485 |
| CoA/Ligand | - | 48 | 96/28 | 96/28 |
| Water | 240 | 280 | 408 | 334 |
| B-factors | | | | |
| Protein | 25.9 | 20.9 | 25.6 | 28.3 |
| CoA/Substrate | - | 33.8 | 48.9/47.6 | 47.5/38.2 |
| Water | 37.0 | 34.6 | 35.5 | 34.5 |
| R.m.s deviations | | | | |
| Bond lengths (Å) | 0.003 | 0.003 | 0.003 | 0.003 |
| Bond angles (°) | 1.060 | 1.066 | 1.064 | 1.064 |

1. Highest resolution shell is shown in parenthesis.
2. R-factor = $\Sigma(|F_{obs}|-k|F_{calc}|)/\Sigma |F_{obs}|$ and R-free is the R value for a test set of reflections consisting of a random 5% of the diffraction data not used in refinement.

**Table S2.** Summary of mutations in spontaneous resistance mutants of *E. coli* WM6242 against flavophos. The mutated DNA regions are shown labeled in bold and underlined font. The frequency of each mutation and whether the genome of the mutant was sequenced or the *phn* operon was amplified by PCR is indicated for each genetic perturbation.

| **Mutation characteristics (frequency observed; genome/PCR)** | **DNA region** |
|---|---|
| A to G mutation in *phnE*<br>(25 %; genome) | CGATCCCGCCCGCGCCGACCATGCCGACGACGGTCGCCGAGCGGACGTTGG**(A to G)**TTCGAAGCG |
| 9 nt insertion (GCCGACCAT) in *phnE*<br>(25 %; genome) | GCGAATCGCTTCCCACAGGGTGACGCCGATCCCGCCCGC**GCCGACCAT**GCCGACCATGCCGACGACGGTCGCCG |
| Large deletion of TOR and Phn transporter components<br>(50 %; genome) | CGTCCAGCCGGTAATTTCAATTTGCAGCCAGTCGCC**/torC-torR-torT-phnE-phnD-phnC-lacI/**CGTTCCGCTATCGGCTGAATTTGATTGCGAGTGAGATATTTATGCCAGCCAGCCAGACGCAGACGCGCCGAGACAGAACTTAATGGG |
| G to A mutation in *phnC*<br>(100%, PCR amplified) | AACGGCGTGCTGCCGAGCGCGCCAATCAGCACGTTCTCCAGTACGCTCAGGCGGTTCACCAGGTTGAATTGTT**(G to A)**GAAGATGTAGCCGGTGTTGGCGCGGCTTTTGCGGATATCGCGCGCCAGACGGCCTTCGCGCTGGACTGTGCGGCCCAGCAGCTCGATATGGCTGCCG |
| 5 nt (CGATG) mutation in *phnE*<br>(75%, PCR amplified) | CCGTTGAGAGAGGAAATCCAGCAGGCTGACCGTGA**CGATG**CGATGATAAGCACCATCAGGGCGCAGGTTTGTTGGAACTGGAAACCGCGAATCGCTTCCCACAGGGTGACGCCGATCCCGC |
| G-insertion in noncoding region between proM tRNA and hisR tRNA<br>(not related to *phn*)<br>(100 %; genome) | TGCGCTACTCGCCGAAATACTGCTTTTTGAATTTTTAGTTCAATTCTTTAAAGTCGTGGTGCGA**G**GGGGGGACTTGAACCCCCACGTCCGTAAGGACACT |

| | |
|---|---|
| C-insertion in noncoding region<br>(not related to *phn*)<br>(100 %; genome) | ACGTATTACGCCCGGCAAGCGACATCAACG<br>TCAGGAT**C**CCCCCTCACCGGCGT |
| G-insertion in noncoding region<br>(not related to *phn*)<br>(100 %; genome) | TTCACGATGAAACCCGGGCGCAAATGGATA<br>AACTATGGTTGGCGCTGGC**G**GGGGGGAAAC<br>GAGAAAATTCACCGACGTT |
| G-insertion in noncoding region<br>(not related to *phn*)<br>(100 %) | TTTGCATCAGGATGACCTCGCGCGATTGTTG<br>GCGTTGAGCGAAACCAGCCGCAC**G**GGGGGG<br>ATCCTCGCCGCACCAGT |
| G-insertion in noncoding region<br>(not related to *phn*)<br>(100 %) | GACGGTTTCCTGTTTTGCTGATAATCCGGAA<br>AATATTCTCGCCTGGTTAGGGCCGGCAATTG<br>GTCCACGCGCGTTCGAAGT**G**GGGGGGGAGG<br>TTCGCGAGGCGTTTATGGCAGTAGACGCTAA |
| A-insertion ahead of *yecD* gene<br>(not related to *phn*)<br>(100 %) | TTAGCATTAAGTTCAAGCATAATGACTCCTG<br>**A**AAAAACAAAATCGTGCCTCACACCTTAAT<br>G |
| G-insertion noncoding region<br>(not related to *phn*)<br>(100 %) | TGACTCTTAACAACGATTCCGCGGCGTATCA<br>GGGTACGACGGATATCGT**G**GGGGGGGAAAT<br>T |
| C-insertion noncoding region again<br>(not related to *phn*)<br>(100 %) | TCTGCCGCCCGCTATCCGGGGCGGCCTTCCC<br>TGCCGATTAG**C**CCCCCCCCTTTCCTCTTTG |

**Table S3.** Primers for plasmid assembly, site-directed mutagenesis, and codon-optimized genes.

| Name | 5' Sequence 3' |
|---|---|
| *Bm*BsfD-MCS2 F | GTATAAGAAGGAGATATACATATGACTAAGCGTAAGACCAT |
| *Bm*BsfD-MCS2 R | GTTTCTTTACCAGACTCGAGGTCACGCAGGCTGCGTCTC |
| *Bs*BsfD-MCS2 F | ATAAGAAGGAGATATACATAATGGCCACGCGCCGTACC |
| *Bs*BsfD-MCS2 R | TTTCTTTACCAGACTCGAGGTTATGCGGCAAGCGCGCG |
| psfC-*Bs*BsfD F | GCGACGCTGAAATGATTAATATGCGTCCGGCGTAGAGG |
| psfC-*Bs*BsfD R | TGCTCAGCGGTGGCAGCAGCCAAAAAACCCCTCAAGACCCG |
| *Bm*BsfE F | ACTTTAATAAGGAGATATACATGCCATTTAGCGTGATC |
| *Bm*BsfE R | GTTCGACTTAAGCATTATGCTTATTTCGTAAGTGCGCC |
| *Bs*BsfD (pET28) F | CTGGTGCCGCGCGGCAGCCATATGGCCACGCGCCGTACC |
| *Bs*BsfD (pET28) R | TGGTGGTGGTGCTCGAGTGCGGCCGCTTATGCGGCAAGCGCGCG |
| *Bm*BsfD (pET15) F | CTGGTGCCGCGCGGCAGCCATATGACTAAGCGTAAGACCATTTTAAC TTGTG |
| *Bm*BsfD (pET15) R | CTTTGTTAGCAGCCGGATCCTCACGCAGGCTGCGTCTC |
| *Bm*BsfB-pBAD F | GGCTAACAGGAGGAATTAACATGAAACGTATCGCCGTAATC |
| *Bm*BsfB-pBAD R | GCCAAAACAGCCAAGCTTCGTCACAACATAGCACATACAG |
| *Bs*RibE F | GGCTAACAGGAGGAATTAACATGGAGATCGGCCAGTATC |
| *Bs*RibE R | GCCAAAACAGCCAAGCTTCGTCATGCACGTTCTTCTTC |
| *Ec*RibE F | GGCTAACAGGAGGAATTAACATGAATATTATCGAAGCGAAC |
| *Ec*RibE R | GCCAAAACAGCCAAGCTTCGTTAGGCTTTAATTGCTTTCAG |
| *Ec*RibC F | GGCTAACAGGAGGAATTAACATGTTTACGGGGATTGTAC |
| *Ec*RibC R | GCCAAAACAGCCAAGCTTCGTCAGGCTTCTGTACCTGG |
| RP_*Bsu*Ls | CTCCTGCTAGCCCAAAAAAA |
| FP_*Bsu*Ls | ATGCAAGCTTGGCTGTTTTG |
| RP_*Aa*Ls | CGAATTCCCATATGGTACCAGCTGCAGATCTCGAGgcgcaggcttttaaacaagtt ggccatctcgatt |
| FP_*As*Ls | ACCCGTTTTTTGGGCTAACAGGAGGAATTAACCATGGatgcaaatatatgaggg aaaactaacagctgaaggcct |
| gBlock for *Bm*BsfD codon-optimization from *B.* | ATGACTAAGCGTAAGACCATTTTAACTTGTGCAGTTACCGGTTCGCAG ACTCGTCAAGATCAAAACCCGAATTTGCCTATCACACCCCGCCAGAT TGCAGAAAGCTCGTTGGAAGCAGCAGATGCAGGTGCTGCCGTGGTTC ATTTACACGTACGTGATCCTCATACTGGTAAAGCTAGCATGGATATCG GACATTATCGTGAAGTTGTGGAGCGTATCCGCGAACGTAATCCTGCG TTAATTTTGAACATCACCACTGGCCCGGGAGGTCGTTTTCAGCCAGGT |

| | |
|---|---|
| *mayonis* sp. BDU6 | GAACCCGACCCTCTGGTCCCGGGACCTCGCACGAATCTTTTACCTCCC<br>GAACGCCGTGTGGCACACTTGGGTGCTCTGCGCCCTGATATTGCGACT<br>GTTGACCTGGACACCATGTTCTTCGGGGGGGACGTCGTAATTAACAC<br>ACCTGCCAGTATTCGCACGATCGCGCGCGCGATCTATGAAACAGGAG<br>CGATTCCCGAACTGGAATTATTCGATAGCGGAAACTTGCATCTGGCA<br>CGTGACTTGTTTGACGACGGGACCTTCCGCCACCCAGCTATTGCATCT<br>CTGATTGTCGGCATCAAGTATGGAATGCCTGCCACCCCAGAAGCTCT<br>GGTGTTTGCTAAGTCATTGCTGCCACCTGGAGTTGAGTGGACTGGCTT<br>TTCAATCGGCCGTCATGCTTTCGCTATGCTTGCGCAGAGCTTCATTTT<br>GGGTGGGCATGTTCGCATTGGTATGGAAGATACAATTTACATCGATA<br>AAGGTAAATTGGCTTCGGGTAATGGTGAGCTGGTGGACCGTGCTAAA<br>TGGATTGTCGAACGCTTAGGGGGGGAATTGGCATCAGCGGAAGAGGC<br>CCGTGTTCAACTTGGGCTGGAGACGCAGCCTGCGTGA |
| gBlock for *Bs*BsfD codon-optimization from *B. stagnalis* | ATGGCCACGCGCCGTACCATTCTGACCTGTGCGGTCACAGGAAGCCA<br>GACGCGCTTAGAACAGAACCCGCATCTGCCAATCACTCCAAAACAGA<br>TCGCCGATGCCTCCCTGGAGGCTGCCGACGCTGGCGCTGCTATTGTGC<br>ATCTGCACGTGCGTGATCCCGAAACTGGCCGTGCTTCAATGGAGTTG<br>GCTCACTATCAGGAGGTTGTGGATCGCATCCGTCAAAAAAACCCTGC<br>ACTTATTGTGAACATTACCACGGGACCTGGGGGACGTTTTCAGCCTG<br>GGGACGTTGATCCCGCTGTAGCGGGCCCCCGCACAAACTTATTACCC<br>CCTGAGCGCCGCGTAGCCCATTTACCCTCATTACGCCCGGACATTGCT<br>ACGTTAGATTTGGACACAATGTTCTTCGGTAGCGAAGTCGTGATTAAC<br>ACACCGCCGACCATCCGTGCCATCGCTCGTGCCATTCATGCATGTGGT<br>GCAGTTCCTGAGTTAGAGCTTTTTGATATCGGCAATCTGCACTTGGCA<br>CGCGATTTATTTGACGAAGGCGTCCTTCGCTTGCCGGCAATCGCCTCA<br>ATTATCGTAGGTATCAAATACGGGATGCCAGCAACGACTGAGGCCAT<br>GACCTTGGCGCGCTCAATGTTGCCACCGGGAGTAGAGTGGACAGGGT<br>TCAGCATCGGTCGCCACGCCTTCCCCATGTTAGCGCAGTCATTCGTAC<br>TGGGCGGACACGTACGTATTGGCATGGAAGATACCATTTACATTGAG<br>AAAGGAAAACTGGCTGCCTCGAACGCTGAGTTAGTAGACAAGGCAA<br>AATGGATCGTCGAGCAACTGGGCGGGGAGCTTGCTAGCGCAGAAGA<br>GGCCCGCGAGCAATTACGCATCGGCGAGCCCCGCGCGCTTGCCGCAT<br>AA |
| gBlock for *Bm*BsfB codon-optimization from *B. mayonis* sp. BDU6 | ATGAAACGTATCGCCGTAATCTACGGGACGTATCATGAGTCAGAAGC<br>AGCACAAATGCGTGACGCGGTACGCGCAGAAGCTCTTGCTGCTGGTG<br>CCGAAATTGTGTACGAAAAGGGCGTCCCCGGCAGTATGGAAAAGCCG<br>CTGGCAACCAAACGCGCTTTGATGGAAGAAAACGTTGATGCGGTGGT<br>AGTGCTTGGAATCATTGAAAAAGGAGAGACTCAACATGGGTTAGTCA<br>TGGCTCAGGCTGTGATCCGTAGCTTGGTAGACCTTCAACTTGAGTTTA<br>TGAAGCCGGTCGGTGTTGGGATCTTGGGTCCAGACATCCAGTCACAT<br>CAAATCCCGCCTCGCTTACGTCTGTATGCTGCTAATGCCTTGAAGGCT<br>GTATGTGCTATGTTGTGA |
| gBlock for *Bs*RibE codon-optimization | ATGGAGATCGGCCAGTATCAGCCCAACTTAGAAGGCGATGGGCTGCG<br>TATCGGAATTGTTCAGTCACGCTTTAATGAGCCAGTTTGTAATGGTCT<br>TGCGGATGCGTGCGTAGAGGAATTAGAACGTCTTGGCGTGTCAGGGG<br>AAGATGTGCTTCTTGTTTCCGTACCAGGTGCATTAGAGATCCCACTGG |

| | |
|---|---|
| from *B. stagnalis* | CCCTTCAGAAACTGGCTGAATCTGGTCAGTTCGATGCGTTGATCGCTC<br>TGGGAGCAGTTATTCGTGGTGAAACCTATCATTTCGAACTGGTTAGTA<br>ATGAGAGTGGTGCTGGTATCACGCGTATTGGATTAGACTTCAATCTGC<br>CAATCGCCAACGCCGTTCTTACGACGGAGACAGACGAACAGGCAGTT<br>GCGCGCATGACGGAGAAAGGGCGTGACGCGGCCCGCGTTGCTGTAG<br>AAATGGCGAACTTAACAATGGCGCTGGACCAGCTTGGCGATGACGAG<br>GACGAGGATGAAGAAGAAGATGAAGATGACGAAGAAGAACGTGCAT<br>GA |
| gBlock for *Ec*RibE codon-optimization from *E. coli* BL21(DE3) | ATGAATATTATCGAAGCGAACGTTGCGACCCCTGATGCGCGCGTTGC<br>AATCACAATCGCCCGTTTCAACAACTTCATCAATGATTCCTTGCTTGA<br>AGGTGCCATTGACGCCTTGAAGCGTATCGGACAGGTTAAGGATGAAA<br>ATATTACTGTAGTATGGGTTCCTGGGGCCTACGAGTTACCTTTAGCCG<br>CAGGCGCTCTTGCAAAAACGGGCAAGTATGATGCAGTGATCGCTCTG<br>GGTACGGTGATCCGCGGAGGCACCGCTCACTTTGAGTATGTCGCGGG<br>CGGTGCCAGTAACGGGTTAGCTCATGTAGCCCAAGACAGCGAGATTC<br>CGGTTGCTTTTGGCGTTTTAACCACCGAGAGTATTGAGCAGGCAATCG<br>AACGTGCTGGCACTAAGGCAGGCAACAAGGGGGCGGAGGCAGCCCT<br>GACAGCACTGGAAATGATTAACGTACTGAAAGCAATTAAAGCCTAA |
| gBlock for *Bm*BsfE codon-optimization from *B. mayonis* sp. BDU6 | ATGAAATATCTGCTGCCGACGGCTGCAGCTGGCTTATTACTGCTGGCA<br>GCCCAACCGGCGATGGCTATGCCATTTAGCGTGATCCTGATCGTTCTG<br>TTTGCAGCCTTGCTGCACGCCACCTGGAACGCTTTAATTCGTTCCAGT<br>GGTTCACGCCTGTGGTCATCCACGGTTCTGTGTATCGCCATGGGTCTG<br>TTTGCGCTCTGCTTTGTGCCTCTTCGCCCGATTCCGCCGCGCGACTCGT<br>GGGCGTACATTGTCGCAAGCGCGGTCTTACACGTCGTGTATAACCTC<br>ATGCTGGTACGTGCTTACCAACGTGGCGAGCTGTCAGTAGCTTACCCC<br>ATTGCACGTGGCACTTCCCCGGTACTCGTTACCACCGGTGCCGCGCTG<br>TTTGCCGGCGAACGCATTGGCGCGCCCACCCTGCTGGGATTACTGCTG<br>ATCTCCGGTGGCATCTTTGCGGTTGGTTTAGATCGTCGGGCGCTTGGC<br>GACGATGCCTGGACGCGCGCATTGCCGGATGCACTGGGCACCGGGGC<br>CTCTATTGCCGCCTATTCAGTGGTGGACGGCATTGGCGCCCGTGCCGC<br>CGGCGATAGCATCGGATATGCGGCCTGGATGTTCGCCTTGACCGCGC<br>TGATGATGCTCCCTACATATTGGCTGTTAGAAGGCCGTCTGACTCTGG<br>GCGATCGCAAGCGCGAACTCGTCAAAGCTGGCTTTGGGGGGATGTCC<br>GCAGCCCTCGGTTATGGTATCGTAATTTGGGCGATGAAACAGGGCGC<br>CATGGGTCCGGTGAGCTCCCTGCGTGAAACAAGCGTTGTATTTGCTGC<br>GCTTATTGGCCGTTTCTTTTTGAGCGAACCCGCGTCGGCTCGCAAAGT<br>ATTAGGCTGCAGTATCATTACGACGGGTACGGTTCTGATTGGGTTCAG<br>TGGCGCACTTACGAAATAA |
| gBlock for *Bs*BsfE codon-optimization from *B. stagnalis* | ATGAAGTATCTTTTACCAACCGCGGCCGCTGGGCTGCTGTTATTGGCG<br>GCGCAGCCTGCAATGGCAATGCCGATTCCCATCCTGTTCGCGGTACTG<br>AGTGCAGCTCTTCTCCATGCTTCATGGAATGCCCTGGTGCGCAGCAGC<br>GGCGATCGTCTGTGGTCTGCGACTGTCCTCTGCATCGCCATGGGCCTG<br>TTTGCCGCGGCCTTAATTCCTTTTGCACCCTTCCCGGTGGCCGCCAGT<br>TGGACCTGTGTGGTGGCGTCGGCGGTGCTCCATGTGCTGTACAACCTG<br>CTGTTGGTGCATAGTTATCATCAAGGAGACTTGAGCGTGAGCTATCC<br>AGTTGCGCGTGGGACGTCCCCGTTGCTTGTTACGCTGGGAGCGGCGG |

| | |
|---|---|
| | TTTTTGCAGGTGAGCGCCTGGGAGCGGGTGCCGTTGCGGGTCTTCTGA<br>TGATCTCGGCGGGGATCGTTGCGATTGGGTTAGATCGGGAAAAAGTT<br>TCTCAGCAACGTGTGACAAAAGCGCTCCCAGCAGCGTTGGCAACCGG<br>CGCTACCATCGCTGCGTATTCCGTGGTTGATGGTATCGGCGTGCGGCA<br>CAATGGGGATGCGACGGGCTATACCGCGTGGATGTTCATGCTGACGG<br>CCTGGATGATGGCCGCATATTTTCGCGTGGTCAAAGGCCAATTCGG<br>TTAAGCGGAAGCCGCGCGGAACTGGCGAAAGCAGGTTTTGGCGGGGT<br>CTTTGCAGCTCTGGCCTACGGTATCGTGATTTGGGCGATGCAGCGCGG<br>CCCAATGGGTCCGATCTCGAGTTTACGGAAACCTCAGTCGTCTTTGC<br>TGCTCTGATCGGTCGTGTCTGTCTGGGTGAACGTTTGTCCATTCGCCG<br>TAGCGCGGGCTGTGGTATTATCGCAGCAGGCGCCGTTCTGATCTCTCT<br>GTCTGGCCTCGCCCGCTAA |
| gBlock for pRSF-Duet_*Ec*RibE | GGGCAGCAGCCATCACCATCATCACCACAGCCAGGATCCATGAATAT<br>TATCGAAGCGAACGTTGCGACCCCTGATGCGCGCGTTGCAATCACAA<br>TCGCCCGTTTCAACAACTTCATCAATGATTCCTTGCTTGAAGGTGCCA<br>TTGACGCCTTGAAGCGTATCGGACAGGTTAAGGATGAAAATATTACT<br>GTAGTATGGGTTCCTGGGGCCTACGAGTTACCTTTAGCCGCAGGCGCT<br>CTTGCAAAAACGGGCAAGTATGATGCAGTGATCGCTCTGGGTACGGT<br>GATCCGCGGAGGCACCGCTCACTTTGAGTATGTCGCGGGCGGTGCCA<br>GTAACGGGTTAGCTCATGTAGCCCAAGACAGCGAGATTCCGGTTGCT<br>TTTGGCGTTTTAACCACCGAGAGTATTGAGCAGGCAATCGAACGTGC<br>TGGCACTAAGGCAGGCAACAAGGGGGCGGAGGCAGCCCTGACAGCA<br>CTGGAAATGATTAACGTACTGAAAGCAATTAAAGCCTAAGAATTCGA<br>GCTCGGCGCGCCTGCAGGTCGACAAGCTTGC |
| gBlock for pRSF-Duet_*Ec*RibB | ATGGGCAGCAGCCATCACCATCATCACCACAGCCAGGATCCaATGAA<br>TCAGACGCTACTTTCCTCTTTTGGTACGCCTTTCGAACGTGTTGAAAA<br>TGCACTGGCTGCGCTGCGTGAAGGACGCGGTGTAATGGTGCTTGATG<br>ATGAAGACCGTGAAAACGAAGGTGATATGATCTTCCCGGCAGAAACC<br>ATGACTGTTGAGCAGATGGCGCTGACCATTCGCCACGGTAGCGGTAT<br>TGTTTGCCTGTGCATTACTGAAGATCGCCGTAAACAACTCGATCTGCC<br>AATGATGGTAGAAAATAACACCAGCGCCTATGGCACCGGTTTTACCG<br>TGACCATTGAAGCAGCTGAAGGTGTGACTACCGGTGTTTCTGCCGCT<br>GACCGTATTACGACCGTTCGCGCAGCGATTGCCGATGGCGCAAAACC<br>GTCAGATCTGAATCGTCCTGGCCACGTTTTCCCACTTCGCGCTCAGGC<br>AGGTGGTGTACTGACGCGTGGCGGTCATACTGAAGCAACTATTGATC<br>TGATGACGCTGGCAGGCTTTAAACCGGCTGGTGTACTGTGTGAGCTG<br>ACTAATGACGATGGCACGATGGCGCGTGCACCAGAGTGTATTGAGTT<br>TGCCAATAAACACAATATGGCGCTCGTGACTATTGAAGACCTGGTGG<br>CATACCGTCAGGCACATGAGCGTAAAGCCAGCTGACGAGCTCGGCGC<br>GCCTGCAGGTCGACAAGCTTGC |
| JRF_His-*Ec*RibE/B_F | GGGCAGCAGCCATCACCATCATCACCACAGCCAGGATCCATGAATAT<br>TATCGAAGCGAAC |
| JRF_His-*Ec*RibE/B_R | GCAAGCTTGTCGACCTGCAGGCGCGCCGAGCTCGAATTCTTAGGCTTT<br>AATTGCTTTCAG |

| | |
|---|---|
| G-block for pBADCDF-BsuLs | TTTTTTTGGGCTAGCAGGAGGAATTCACCATGAACATTATCCAGGGCAATTTAGTTGGCACAGGCCTTAAAATCGGGATTGTGGTAGGACGCTTCAATGACTTCATTACCAGCAAACTTCTGTCCGGCGCAGAAGATGCCCTGCTTCGTCACGGTGTGGATACTAACGACATTGACGTGGCTTGGGTTCCGGGCGCATTTGAAATTCCATTTGCCGCGAAAAAAATGGCGGAAACGAAGAAATATGATGCGATTATTACTCTTGGCACGGTAATTCGCGGTGCGACGACGCATTATGATTACGTCTGCAACGAGGCCGCCAAAGGTATCGCGCAGGCAGCGAACACAACCGGCGTGCCGGTAATCTTCGGCATTGTTACTACTGAGAATATCGAACAAGCTATTGAACGTGCGGGTACCAAAGCGGGAAATAAGGGAGTGGACTGCGCCGTGAGCGCAATCGAAATGGCAAACTTAAACCGTAGTTTTGAATAAGTACCCGGGGATCCTCTAGAGTCGACCTGCAGGCATGCAAGCTTGGCTGTTTTG |
| G-block for pBADCDF-BsuLs_ H88K_K131N | TTTTTTTGGGCTAGCAGGAGGAATTCACCATGAACATTATCCAGGGCAATTTAGTTGGCACAGGCCTTAAAATCGGGATTGTGGTAGGACGCTTCAATGACTTCATTACCAGCAAACTTCTGTCCGGCGCAGAAGATGCCCTGCTTCGTCACGGTGTGGATACTAACGACATTGACGTGGCTTGGGTTCCGGGCGCATTTGAAATTCCATTTGCCGCGAAAAAAATGGCGGAAACGAAGAAATATGATGCGATTATTACTCTTGGCACGGTAATTCGCGGTGCGACGACGAAATATGATTACGTCTGCAACGAGGCCGCCAAAGGTATCGCGCAGGCAGCGAACACAACCGGCGTGCCGGTAATCTTCGGCATTGTTACTACTGAGAATATCGAACAAGCTATTGAACGTGCGGGTACCAACGCGGGAAATAAGGGAGTGGACTGCGCCGTGAGCGCAATCGAAATGGCAAACTTAAACCGTAGTTTTGAATAAGTACCCGGGGATCCTCTAGAGTCGACCTGCAGGCATGCAAGCTTGGCTGTTTTG |
| G-block for pBADCDF-BsuLs_R127T | TTTTTTTGGGCTAGCAGGAGGAATTCACCATGAACATTATCCAGGGCAATTTAGTTGGCACAGGCCTTAAAATCGGGATTGTGGTAGGACGCTTCAATGACTTCATTACCAGCAAACTTCTGTCCGGCGCAGAAGATGCCCTGCTTCGTCACGGTGTGGATACTAACGACATTGACGTGGCTTGGGTTCCGGGCGCATTTGAAATTCCATTTGCCGCGAAAAAAATGGCGGAAACGAAGAAATATGATGCGATTATTACTCTTGGCACGGTAATTCGCGGTGCGACGACGCATTATGATTACGTCTGCAACGAGGCCGCCAAAGGTATCGCGCAGGCAGCGAACACAACCGGCGTGCCGGTAATCTTCGGCATTGTTACTACTGAGAATATCGAACAAGCTATTGAAACCGCGGGTACCAAAGCGGGAAATAAGGGAGTGGACTGCGCCGTGAGCGCAATCGAAATGGCAAACTTAAACCGTAGTTTTGAATAAGTACCCGGGGATCCTCTAGAGTCGACCTGCAGGCATGCAAGCTTGGCTGTTTTG |
| gBlock for F166Y-*Bs*BsfD | CGGTAGCGAAGTCGTGATTAACACACCGCCGACCATCCGTGCCATCGCTCGTGCCATTCATGCATGTGGTGCAGTTCCTGAGTTAGAGCTTtacGATATCGGCAATCTGCACTTGGCACGCGATTTATTTGACGAAGGCGTCCTTCGCTTGCCGGCAATCGCCTCAATTATCGTAGGTATC |
| *Bs*BsfD F | GCCGGCAATCGCCTCAATTATC |
| *Bs*BsfD R | GAATGGCACGAGCGATGGC |

**Table S4.** Plasmids and microorganisms used in this study.

| Strain or plasmid | Relevant characteristics | Source or reference |
|---|---|---|
| *Escherichia coli* NEB5α (plasmid maintenance) | *fhuA2 Δ(argF-lacZ)U169 phoA glnV44 Φ80 Δ(lacZ)M15 gyrA96 recA1 relA1 endA1 thi-1 hsdR17* | New England BioLabs (Ipswich, MA) |
| *Escherichia coli* BL21(DE3) (protein overexpression) | | New England BioLabs (Ipswich, MA) |
| *Escherichia coli* WM6242 | Phosphonate uptake strain containing IPTG-inducible *phn* operon duplication under $P_{tac}$ control | Eliot et al.[6] |
| *Escherichia coli* WM6242 λDE3 | Phosphonate uptake strain lysogenized with DE3 lysogen for expression of T7 RNA polymerase | This study |
| *Burkholderia* sp. Bp9002 | Strain containing the flavophos BGC | Northern Arizona University |
| *Burkholderia* sp. Bp9097 | Strain containing the flavophos BGC | Northern Arizona University |
| *Burkholderia mayonis* sp. BDU6 | Strain containing the flavophos BGC | Northern Arizona University |
| *Burkholderia* sp. MSMB2051 | Strain containing the flavophos BGC | Northern Arizona University |
| pETDuet | $Amp^R$ empty expression plasmid | Novagen |
| pCDFDuet | $Sm^R$ empty expression plasmid | Novagen |
| pBAD-HisA | AmpR empty expression plasmid with $P_{bad}$ promoter | Thermo Fisher |
| pETDuet_psfC_*Bs*BsfD | $Amp^R$ co-expression of psfC and *Bs*BsfD | This study |
| pETDuet_psfC_*Bm*BsfD | $Amp^R$ co-expression of psfC and *Bm*BsfD | This study |
| pCDFDuet_*Bm*BsfE | $Sm^R$ co-expressions of BsfE from *B.* sp. BDU6 | This study |
| pRSF_psfABC_*Bs*BsfD | $Kan^R$ co-expression of psfABC and *Bs*BsfD | This study |
| pET15b_His-*Bm*BsfD | $Amp^R$ expression of *Bm*BsfD for purification | This study |
| pET28a_His-*Bs*BsfD | $Kan^R$ expression of *Bs*BsfD for purification | This study |
| pBAD_*Bm*BsfB | $Amp^R$ expression of *B. mayonis* sp. BDU6 *bsfB* under arabinose control | This study |
| pBAD_*Bs*RibE | $Amp^R$ expression of *B. stagnalis* lumazine synthase under arabinose control | This study |

| | | |
|---|---|---|
| pBAD_*Ec*RibE | Amp$^R$ expression of *E. coli* lumazine synthase under arabinose control | This study |
| pBAD_*Ec*RibC | Amp$^R$ expression of *E. coli* riboflavin synthase under arabinose control | This study |
